# A Constitutively Closed Pannexin1 Channel in Lipid Bilayer Nanodiscs Assembles as a Large-Pore Heptamer

**DOI:** 10.1101/2020.12.31.425019

**Authors:** Xueyao Jin, Susan A. Leonhardt, Yu-Hsin Chiu, Michael D. Purdy, William E. McIntire, Brad C. Bennett, Douglas A. Bayliss, Mark Yeager

**Author notes:** These authors contributed equally to this work (Xueyao Jin, Susan A. Leonhardt). Howard College of Arts and Sciences, Department of Biological and Environmental Sciences, Samford University, Birmingham, Alabama 35229 (Brad C. Bennett). Institute of Biotechnology and Department of Medical Science, National Tsing Hua University, Hsinchu, Taiwan (Yu-Hsin Chiu). **Corresponding author:** Mark Yeager, M.D., Ph.D., University of Virginia School of Medicine, Department of Molecular Physiology and Biological Physics, Sheridan G. Snyder Translational Research Building, Rm 320, 480 Ray C. Hunt Drive, Charlottesville, VA 22908.

## Abstract

Pannexin 1 (Panx1) channels are widely expressed and play important roles in apoptotic cell clearance, inflammation, blood pressure regulation, neurological disorders, opiate withdrawal, and cancer progression and metastasis. We performed (1) physicochemical analysis on a constitutively closed Panx1 channel (designated fPanx1ΔC) to examine the entire population of particles to detect multiple oligomeric states and (2) cryoEM in the membrane mimetics amphipol A8-35 and lipid bilayer nanodiscs. Our results reveal that the dominant if not exclusive oligomeric state of fPanx1ΔC is a heptamer, in solution and by cryoEM. The Panx1 heptamer provides further structural diversity within the family of large-pore channels, including hexameric LRRC8 (SWELL1) channels and connexin hemichannels, octameric CALHM1 channels and innexin hemichannels, and undecameric CALHM2 channels. Conserved structural themes are a large cytoplasmic vestibule with a diameter that corresponds roughly with the oligomeric state and a 4-helix bundle protomer, albeit with noncanonical helical packing for CALHM1 and CALHM2.

**In Brief:** The 4-helix bundle protomer of a constitutively closed pannexin1 channel assembles as a heptamer in solution and by cryoEM.

## INTRODUCTION

Pannexins (Panxs) were discovered in 2000 on the basis of their limited sequence homology to innexins (Inxs), the invertebrate gap junction channels, and are a distinct channel subfamily within the mammalian genome (Panchin et al., 2000). Despite a lack of sequence similarity to connexins (Cxs), the vertebrate gap junction channels, Panxs share a common 4 transmembrane (TM) domain topology with Cxs and Inxs, as well as recently identified calcium homeostasis modulator channels (CALHM1 and 2) and the leucine-rich repeat-containing 8 (LRRC8, SWELL) channels (Abascal and Zardoya, 2012; Baranova et al., 2004; Ma et al., 2012; Scemes et al., 2009). Each protein subunit contains TM domains that are connected by two extracellular loops (EL) and one cytoplasmic loop (CL), with the amino (NT) and carboxyl (CT) termini located on the cytoplasmic side (**Fig. 1**). Cxs, Inxs, and CALHM2 can assemble as single membrane hemichannels and can also dock end-to-end to form intercellular gap junction channels. In contrast, Cx and Inx hemichannels, Panxs, LRRC8 (SWELL), and CALHM1 only form single membrane channels that transport molecules between the cytosolic and extracellular space (Dahl and Keane, 2012; Dahl and Locovei, 2006; Sosinsky et al., 2011).

**Figure 1.**
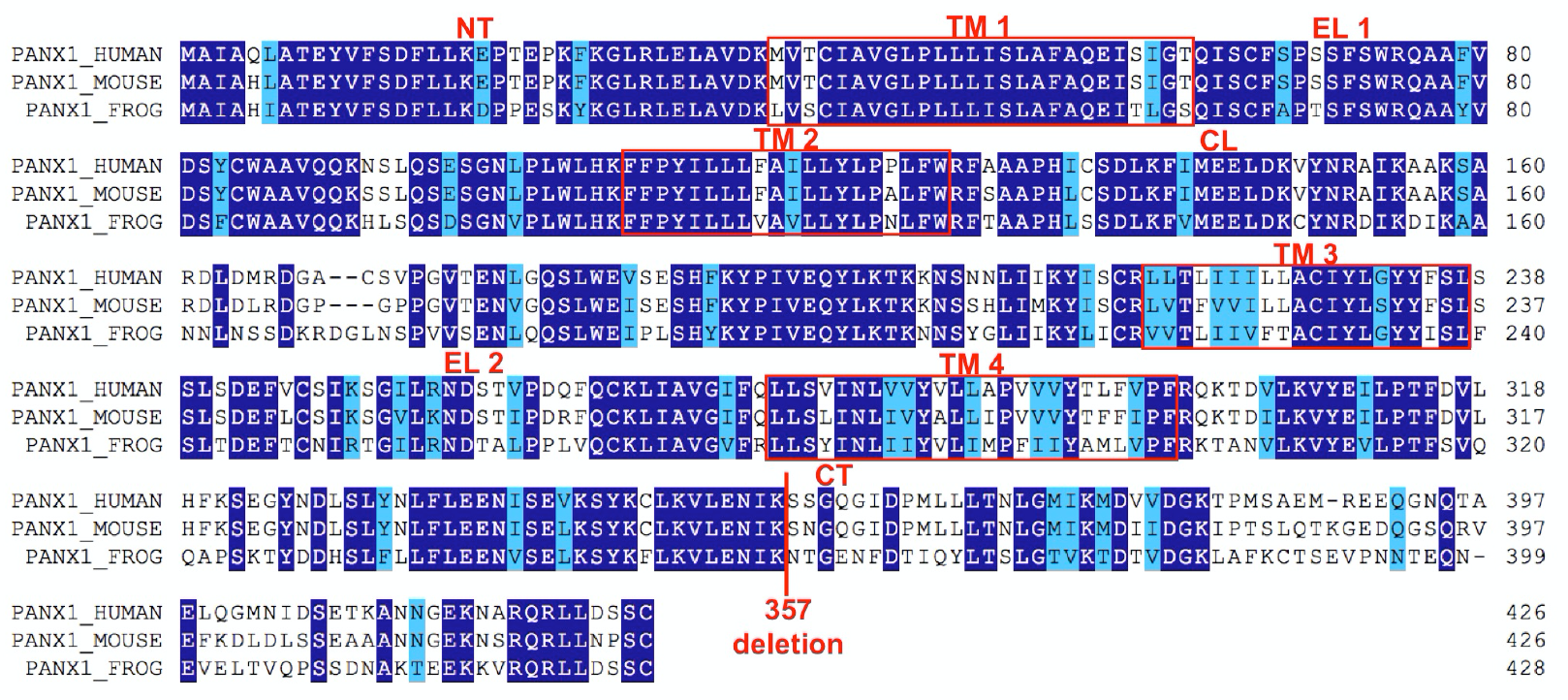
Sequence alignment of human, mouse and frog Panx1. Selected Panx1 isoforms aligned with their amino acid sequence by ClustalW (Thompson et al., 1994). Identical residues and similar residues are colored dark blue and light blue, respectively, using the Multiple Align Show web-based program. fPanx1ΔC resulted from C-terminal truncation at Lys357. TM1-TM4, transmembrane α-helices; EL1 and EL2, extracellular loops; CL, cytoplasmic loop; NT and CT, amino and carboxyl termini. Note the lack of sequence conservation beyond residue 357.

Panx1, one of the three mammalian Panx family members (Panx1-3), is a widely expressed, plasma membrane channel (Baranova et al., 2004). Panx1 is reportedly activated by a variety of physiological conditions, including mechanical stress (Bao et al., 2004; Locovei et al., 2006a), membrane depolarization (Bao et al., 2004; Bruzzone et al., 2003; Ma et al., 2009), increased extracellular potassium (Qiu et al., 2011; Santiago et al., 2011; Silverman et al., 2009; Suadicani et al., 2012; Wang et al., 2014; Wang et al., 2018), increased intracellular calcium (Locovei et al., 2006b; Seminario-Vidal et al., 2009), receptor-mediated signaling pathways (Adamson et al., 2015; Billaud et al., 2015; Billaud et al., 2011; Iglesias et al., 2008; Locovei et al., 2006b; Pelegrin and Surprenant, 2006; Thompson et al., 2008), and caspase-mediated cleavage of the distal C-terminus (Chekeni et al., 2010; Qu et al., 2011; Sandilos et al., 2012; Yang et al., 2015). Upon activation, Panx1 channels allow TM flux of ions, and larger molecules such as nucleotides (ATP, UTP), metabolites and fluorescent dyes (cationic To-Pro, anionic Lucifer Yellow) (Bao et al., 2004; Chekeni et al., 2010; Dolmatova et al., 2012; Medina et al., 2020). Panx1-dependent ATP release plays an important role in diverse physiological and pathological processes, including blood pressure regulation (Billaud et al., 2015; Billaud et al., 2011; Locovei et al., 2006a), apoptotic cell clearance (Chekeni et al., 2010; Poon et al., 2014), inflammation (Kanneganti et al., 2007; Pelegrin and Surprenant, 2006), neurological disorders (Gulbransen et al., 2012; Karatas et al., 2013; Santiago et al., 2011; Thompson et al., 2008; Thompson et al., 2006), opiate withdrawal (Burma et al., 2017) and cancer metastasis (Furlow et al., 2015; Penuela et al., 2012).

Structures of Cx gap junction channels have been studied for decades, and subnanometer and high-resolution structures 2D electron cryocrystallography (Oshima et al., 2007; Unger et al., 1999), single-particle electron cryomicroscopy (cryoEM) (Flores et al., 2020; Khan et al., 2020; Myers et al., 2018) and X-ray crystallography (Bennett et al., 2016; Maeda et al., 2009). Extensive structural studies on Cx gap junction channels have led to a common view that a dodecameric Cx gap junction channel is formed by the docking of two hexameric hemichannels (Bennett et al., 2016; Maeda et al., 2009; Oshima et al., 2007; Unger et al., 1999; Unwin and Ennis, 1984). In a similar fashion Inx-6 octameric hemichannels can also form gap junction channels (Oshima et al., 2016b).

LRRC8A (SWELL1), a volume-regulated anion channel (VRAC), shares the 4 TM domain topology and a common ancestral gene with Panx1 (Abascal and Zardoya, 2012). In 2018, the high-resolution structure of LRRC8A (SWELL1) was determined by a combination of X-ray crystallography and single-particle cryoEM (Deneka et al., 2018). Within months, three additional cryoEM structures appeared (Kasuya et al., 2018; Kefauver et al., 2018; Kern et al., 2019). All structures demonstrated that LRRC8 assembles as a hexamer.

Over the last ∼15 years Panxs were considered to be analogous to Cxs, in that they share a common 4 TM domain membrane topology and form similar large-pore channels (Baranova et al., 2004; Boassa et al., 2007; Bruzzone et al., 2003; Locovei et al., 2006a; Locovei et al., 2006b; Penuela et al., 2007). A notable difference is that pannexins do not form gap junction channels except when the extracellular carbohydrate moiety is removed (PDB code: 6WBN) (Ruan et al., 2020). Analysis of the subunit stoichiometry by electron microscopy (EM), chemical crosslinking and single-molecule photobleaching suggested that Panx1 was a hexamer (Ambrosi et al., 2010; Boassa et al., 2007; Chiu et al., 2017) analogous to Cx hemichannels. Hence, the recent observations that Panx1 assembles as a heptamer were unexpected.

Similar to the impressive productivity in the structural biology of LRRC8A (SWELL1) channels, six publications appeared in 2020 describing 16 independent high-resolution, 3D cryoEM reconstructions of various constructs of frog and human Panx1 channels: PDB codes 6UZY, 6V6D (Deng et al., 2020), PDB codes 6M66, 6M67, 6M68 (Jin et al., 2020), PDB code 6VD7 (Michalski et al., 2020), PDB codes 6LTN, 6LTO (Mou et al., 2020), PDB code 6MO2 (Qu et al., 2020), PDB codes 6WBF, 6WBG, 6WBI, 6WBK, 6WBL, 6WBM, 6WBN (Ruan et al., 2020). All structures demonstrate that Panx1 assembles as a heptamer.

To further explore the structure and stoichiometry of Panx1 channels, we examined frog Panx1 that we truncated at Lys357 due to the low conservation of sequence thereafter (designated fPanx1ΔC) (**Fig. 1**). Otherwise, the sequences of frog and human Panx1 are highly similar (**Fig. 1**). A further advantage of this construct is that electrophysiology demonstrated that fPanx1ΔC is basally closed and presumably exists in a single conformation. Furthermore, fPanx1ΔC could be activated by α1D adrenoceptor stimulation as has been shown for human Panx1. We note that fPanx1ΔC is truncated upstream of the C-terminal caspase site, and thus the receptor activation process is independent of quantized C-terminal cleavage-activation (Chiu et al., 2017). The expression level of fPanx1ΔC was also higher than human Panx1. In this study we performed physicochemical and cryoEM analysis to examine whether fPanx1ΔC exists in multiple oligomeric states. Our results indicate that the dominant if not exclusive oligomeric state of fPanx1ΔC is a heptamer in solution and by cryoEM.

## RESULTS

### Functional validation of fPanx1ΔC

Whole-cell patch clamp recordings obtained from HEK293 cells transfected with fPanx1ΔC showed that unstimulated channels were basally silent (**Fig. 2**), similar to earlier reports of basally silent, full-length human Panx1 channels (Chekeni et al., 2010; Chiu et al., 2017). Panx1 channels can be activated by G protein-coupled receptor (GPCR) mediated signaling pathways (Billaud et al., 2011; Blum et al., 2008; Gödecke et al., 2012; Locovei et al., 2006b; Seminario-Vidal et al., 2009), such as phenylephrine binding to α1 adrenergic receptors, which triggers ATP release from activated Panx1 channels to regulate blood pressure (Billaud et al., 2015; Billaud et al., 2011; Billaud et al., 2012). To test whether fPanx1ΔC was functional via GPCR-stimulation, we co-transfected α1D adrenoceptors (α1DRs) and fPanx1ΔC in HEK293 cells (**Fig. 2a**). Phenylephrine (PE) induced an outwardly rectifying current, which was abolished by carbenoxolone (CBX), a known Panx1 inhibitor (**Fig. 2b**). These results suggest that fPanx1ΔC channels, like the human ortholog, are in a basally closed state that can be activated by α1DR stimulation. Because the fPanx1ΔC channel is in a closed conformation, despite the C-terminal truncation upstream of the caspase cleavage site, there may be an activation gate located elsewhere in the structure.

**Figure 2.**
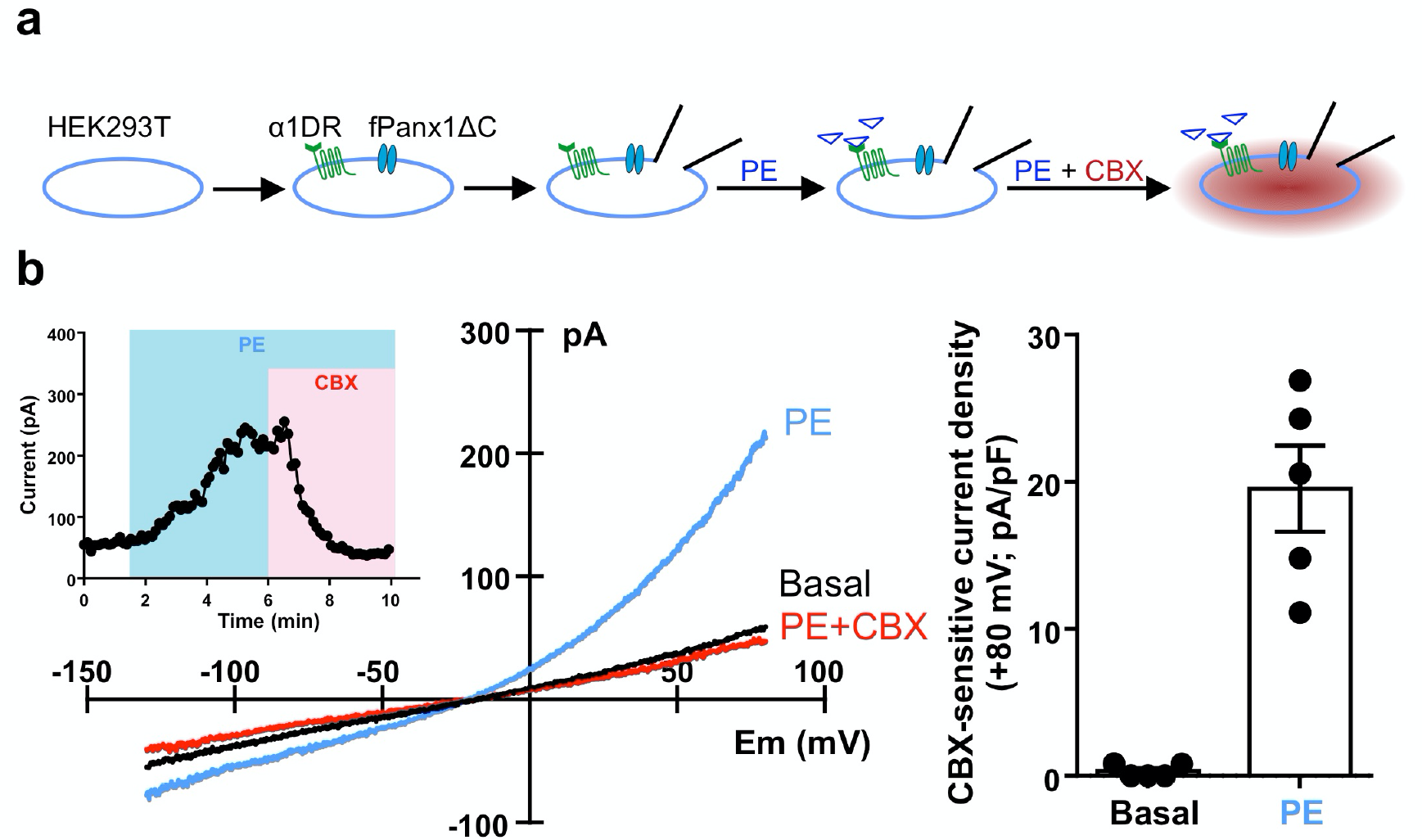
fPanx1ΔC channels are in a basally closed conformation and can be activated by α1D adrenoceptor stimulation. (**a**) Schematic showing the whole-cell recording procedure. (**b**) Whole-cell recordings from HEK293T cells co-expressing α1D adrenoceptors (α1DRs) and fPanx1ΔC channels. fPanx1ΔC channel activity was induced by phenylephrine (PE, 20 µM), and abolished by carbenoxolone (CBX, 50 µM).

### Expression, purification and physicochemical characterization of fPanx1ΔC

Table S1 summarizes the constructs, expression systems and conditions for solubilization and purification. Best results were obtained using fPanx1ΔC (residues 1-357) with a thrombin cleavage site and Strep tag at the C-terminus. fPanx1ΔC was expressed in *Sf9* insect cells, extracted in the detergent n-dodecyl-β-D-maltopyranoside containing cholesteryl hemisuccinate (DDM/CHS) and purified using strep-tactin affinity chromatography. SDS-PAGE and immunoblot analysis showed that the fPanx1ΔC monomer band migrated at ∼37 kDa, which is smaller than the calculated mass of 43 kDa (**Fig. 3a**). It was proposed that Panx1 subunits form a hexamer by examination of DSP (dithiobis(succinimidyl propionate)) crosslinked rat Panx1(Boassa et al., 2007). Here, we used glutaraldehyde to crosslink fPanx1ΔC in DDM/CHS and observed a protein band at ∼250 kDa by SDS-PAGE (**Fig. 3a**).

**Figure 3.**
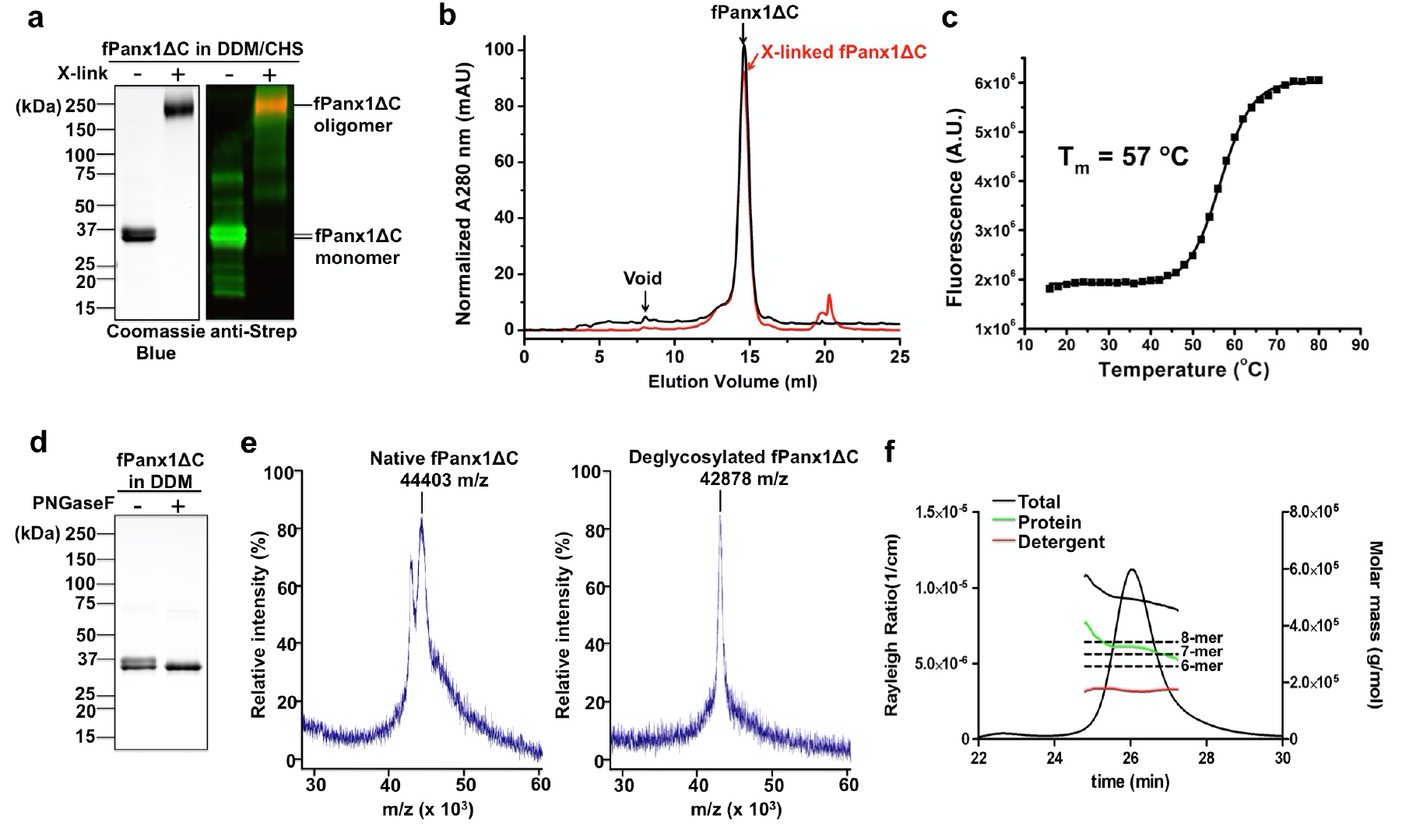
Physicochemical analysis of fPanx1ΔC. (**a**) Coomassie Blue stained SDS-PAGE and immunoblotting probed with anti-Strep antibodies showed monomeric fPanx1ΔC, and oligomeric fPanx1ΔC crosslinked by 0.1% glutaradehyde. (**b**) SEC showed both native (black) and crosslinked (red) fPanx1ΔC were homogenous, with minimal aggregation (void) and migrated at the same elution volume. (**c**) Fluorescence thermostability assay showed that fPanx1ΔC was thermally stable, with a melting temperature (T_m_) of 57 °C. (**d**) Coomassie Blue stained SDS-PAGE showed fPanx1ΔC was deglycosylated with PNGaseF treatment. (**e**) MALDI-MS showed mass-to-charge (m/z) ratio of native (44.403 kDa) and deglycosylated (42.878 kDa) fPanx1ΔC by PNGaseF. (**f**) SEC-MALS showed that the mass of fPanx1ΔC was 320 ± 16 kDa, with hexameric, heptameric and octameric oligomers having masses of 258, 301 and 344 kDa, respectively. (**a-c**) fPanx1ΔC purified in DDM/CHS; **(d-f)** fPanx1ΔC purified in DDM.

Analytical SEC of non-crosslinked fPanx1ΔC showed a single absorbance peak with minimal aggregation and the same elution volume as crosslinked fPanx1ΔC, indicating that fPanx1ΔC forms a stable oligomer (**Fig. 3b**). The thermal stability of fPanx1ΔC was assessed by a fluorescence-based thermal stability assay, in which the quantum yield increases upon temperature-induced protein unfolding, allowing cysteine residues embedded within the protein interior to become accessible to a fluorophore (Alexandrov et al., 2008). fPanx1ΔC started to unfold at 50 °C, and the estimated melting temperature (T_m_) was 57 °C, indicating that fPanx1ΔC is thermally stable in the detergent DDM/CHS (**Fig. 3c**).

Rat Panx1 expressed in HEK293T cells can be N-glycosylated at Asn254, resulting in three protein bands that represent the non-glycosylated core protein (lower band), the high mannose-type glycoprotein (intermediate band), and the fully processed mature-type glycoprotein (upper band), respectively (Boassa et al., 2007). For fPanx1ΔC expressed in *Sf9* cells, we observed two fPanx1ΔC protein bands by SDS-PAGE in DDM/CHS (**Fig. 3a,d**), amphipol A8-35 **(Fig. 4a)** and in nanodiscs **(Fig. 5a)**. As in previous studies (Boassa et al., 2007) the top band of fPanx1ΔC was sensitive to the deglycosylase enzyme PNGaseF (**Fig. 3d**), indicating that fPanx1ΔC is glycosylated in insect cells, albeit likely by a different mechanism relative to mammalian cells. This glycosylation was also confirmed by matrix-assisted laser desorption ionization mass spectrometry (MALDI-MS) of fPanx1ΔC proteins with or without PNGaseF treatment. The molecular weight of deglycosylated fPanx1ΔC (42.878 kDa) agrees closely with the calculated mass value (43.015 kDa) while the larger molecular weight of native fPanx1ΔC (44.403 kDa) could be due to glycosylation in insect cells (**Fig. 3e**).

**Figure 4.**
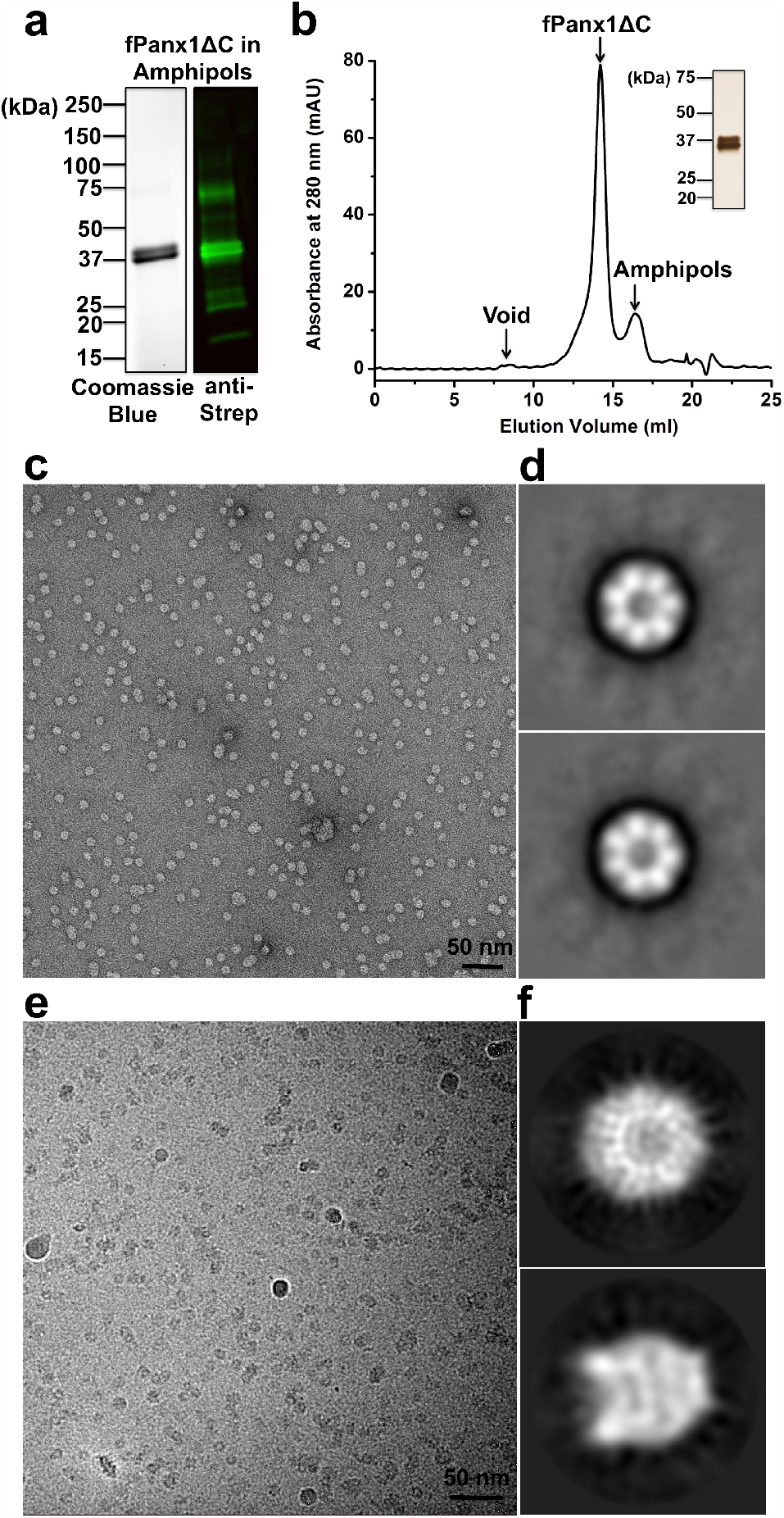
Characterization of fPanx1ΔC in amphipol A8-35 revealed a heptameric channel. (**a**) Coomassie Blue stained SDS-PAGE and immunoblotting probed with anti-Strep antibodies showed monomeric fPanx1ΔC in amphipols. (**b**) SEC showed that fPanx1ΔC was homogenous, with minimal aggregation (Void peak) in amphipol A8-35. fPanx1ΔC protein, amphipol and void peaks are indicated. Inset, silver stained SDS-PAGE of SEC fractions collected from fPanx1ΔC protein peak. (**c**) Electron micrograph of negatively-stained fPanx1ΔC in amphipol A8-35. Scale bar, 50 nm. (**d**) Representative 2D class averages of negatively-stained fPanx1ΔC particles from (**c**) with 1,649 (top) and 1,903 (bottom) particles. The particle box dimension is 357 Å. (**e**) Representative electron cryomicrograph of frozen-hydrated fPanx1ΔC in amphipols. (**f**) 2D class averages from 3,318 particles for the top view (top) and 1,762 particles for the side view (bottom), respectively.

**Figure 5.**
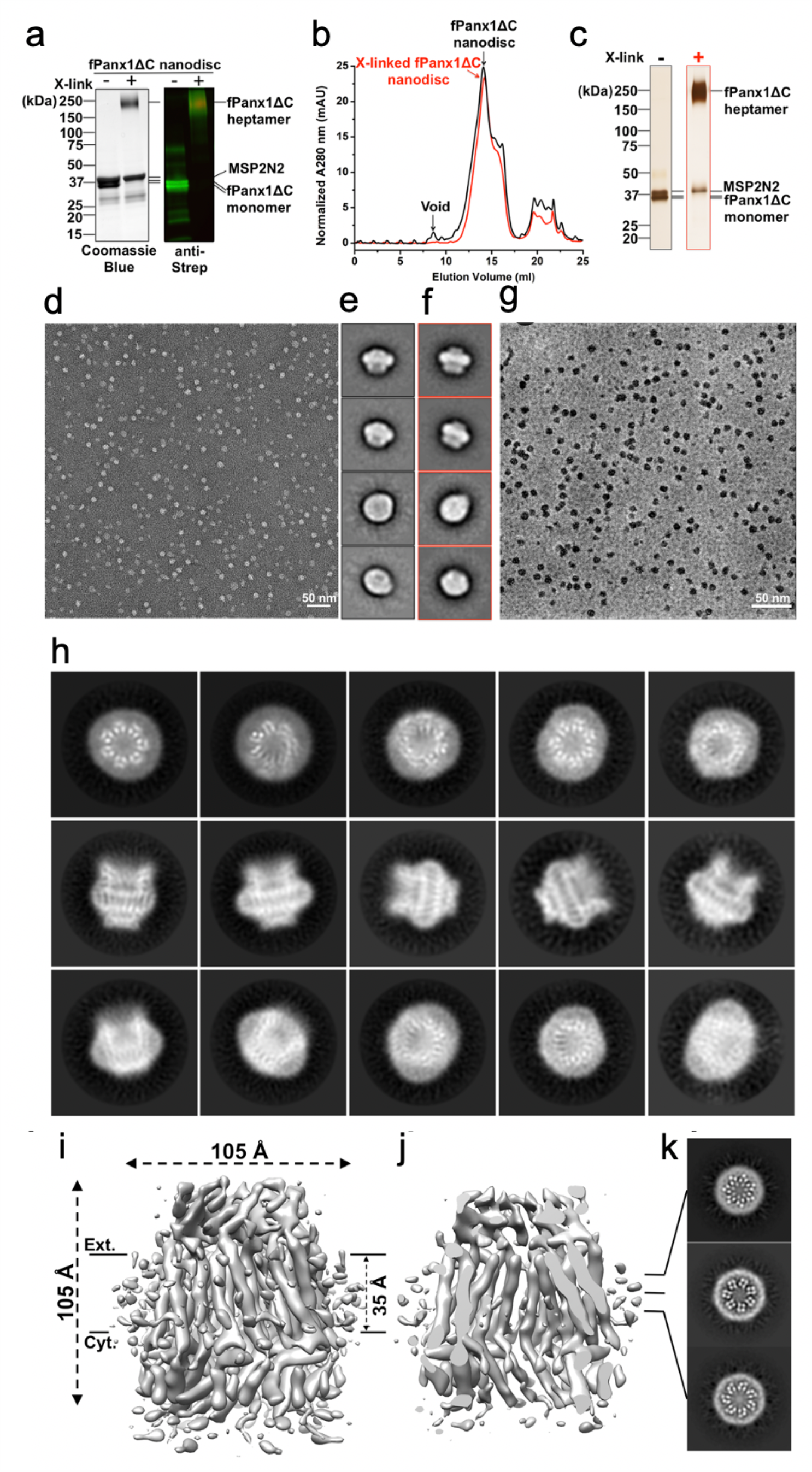
Biochemical characterization and cryoEM of fPanx1ΔC reconstituted in lipid bilayer nanodiscs. (**a**) Coomassie Blue stained SDS-PAGE and immunoblotting probed with anti-Strep antibodies showed monomeric and cross-linked fPanx1ΔC in lipid nanodiscs. (**b**) SEC showed that both native (black) and crosslinked (red) fPanx1ΔC in lipid nanodiscs were homogenous, with minimal aggregation (void peak) and had coincident elution volumes. fPanx1ΔC-nanodisc and void peaks are indicated. (**c**) Silver stained SDS-PAGE of SEC fractions collected from lipid nanodisc embedded-fPanx1ΔC, or crosslinked fPanx1ΔC peaks. (**d**) Electron micrograph of negatively-stained fPanx1ΔC in lipid nanodiscs. Scale bar, 50 nm. (**e**) Representative 2D class averages of negatively-stained fPanx1ΔC in lipid nanodiscs, with 826, 957, 920, and 440 particles (from top to bottom). (**f**) Representative 2D class averages of negatively-stained, crosslinked fPanx1ΔC in lipid nanodiscs, with 806, 517, 324, and 476 particles (from top to bottom). **(g)** Representative electron cryomicrograph of frozen-hydrated fPanx1ΔC in lipid nanodiscs recording using a Volta phase plate. Scale bar, 50 nm. (**h**) 2D class averages of frozen-hydrated fPanx1ΔC in lipid nanodiscs, showing top, side and oblique views. Box dimension, 238 Å. (**i**) Surface and (**j**) cross-sectional side views of the fPanx1ΔC cryoEM density map showing rod-like TM α-helices. Channel height and width were both ∼105 Å. The membrane region defined by the nanodisc was ∼35 Å. Cyt, cytoplasm; Ext, extracellular space. (**k**) Slices through the 3D reconstruction at locations indicated in (**j**) display a heptamer with four circular densities per subunit, ascribed to TM α-helices.

Given the apparent monomer molecular weight of ∼35 kDa by SDS-PAGE, the ∼250 kDa oligomer would correspond to a heptamer. However, it is difficult to determine an accurate oligomeric state just by the molecular weight estimated *via* SDS-PAGE. Thus, we performed size exclusion chromatography combined with multi-angle light scattering (SEC-MALS) (**Fig. 3f**). SEC-MALS yielded an averaged molar mass of fPanx1ΔC at 320.1 kDa. Given a systematic error of 5% (∼16 kDa), the observed mass is between that expected for a heptamer (43.015 kDa x 7 = 301.105 kDa) and an octamer (43.015 kDa x 8 = 344.120 kDa), as calculated from the amino acid sequence.

### EM analysis of fPanx1ΔC channels in the membrane mimetic amphipol A8-35

Upon exchange of fPanx1C into amphipol A8-35, SEC showed that there was almost no aggregation as indicated by a minimal void peak (**Fig. S1b**). SEC was also useful for removing free amphipols. EM of negatively-stained and frozen-hydrated fPanx1ΔC in amphipols showed homogeneous and monodisperse particles (**Fig. 4c,e, Table S2**), and 2D class averages without imposing symmetry displayed a heptamer **(Fig. 4d)**. To verify the oligomeric state of fPanx1ΔC, we performed cryoEM, and individual particles had good contrast and were readily identified in the cryomicrographs (**Fig. 4e**). Interestingly, addition of Ca^2+^ improved the homogeneity and symmetry of the particles (**Fig. S1**). 2D class averages of top view images with no enforced symmetry showed that fPanx1ΔC forms a ring-shaped heptamer (**Fig. 4f**).

### EM analysis of fPanx1ΔC channels in lipid bilayer nanodiscs

We also reconstituted fPanx1ΔC into nanodiscs assembled from soybean polar lipids and the Membrane Scaffold Protein 2N2 (MSP2N2) (**Fig. 5, S2 and S3**). We selected MSP2N2 because it forms nanodiscs of 150-165 Å in diameter, which would be expected to accommodate hexameric Cx26 (Maeda et al., 2009) and octameric Inx-6 (Oshima et al., 2016b) hemichannels, with diameters of 92 Å and 110 Å, respectively. fPanx1ΔC was extracted and purified in DDM/CHS, and then reconstituted into lipid nanodiscs at a molar ratio of 1:1.25:125 (fPanx1ΔC:MSP2N2:lipids). Negative-stain EM showed monodisperse fPanx1ΔC-nanodisc particles (**Fig. 5d)** with different orientations, including side views of the channel with a band-like density contributed by the lipid bilayer and scaffold protein (**Fig. 5d,e, Table S2**). CryoEM was then performed using a Titan Krios electron microscope operating at 300 kV and equipped with a Gatan K2 Summit direct electron detector, energy filter and a Volta phase plate (VPP) at low defocus (−0.5 μm) **(Table S3)**. The VPP greatly improved contrast of the particles, which were homogenous and monodisperse **(Fig. 5g)**. An initial particle data set of 687,949 particles was subjected to reference-free 2D classification, and top view class averages verified a heptameric oligomeric state (**Fig. 5h**). Side view class averages displayed vertical “striping” that is characteristic for TM α-helices. A subset of 319,685 particles derived from 2D class averages was subjected to 3D classification, in which Classes 6 and Class 8 of the ten classes contained features consistent with TM α-helices **(Fig. S3a)**. Class 6 displayed a larger diameter nanodisc compared with Class 8. (Large and small nanodiscs were also observed for the MsbA-nanodisc complex (Mi et al., 2017).) The fPanx1ΔC small-nanodisc class contained a sufficient distribution of particle orientations with top and side views **(Fig. S3b)**, which were used for further 3D refinement (**Fig. S3c,d**). A 3D map was reconstructed from 38,724 particles within the small-nanodisc class, in which C7 symmetry was imposed for 3D auto-refinement. Estimation of the local resolution showed a narrow range (6.6 Å to 7.6 Å), in which the TM region has slightly higher resolution than the cytoplasmic and extracellular regions (**Fig. S3c**). On the basis of the gold-standard Fourier Shell Correlation (FSC) criterion of 0.143, the overall resolution was ∼7.0 Å (**Fig. S3d**). **Table S2** summarizes the data collection and image processing statistics.

The fPanx1ΔC channel comprises three domains: cytoplasmic, transmembrane (TM), and extracellular (**Fig. 5i,j; Fig. 6**). The TM domain is approximately perpendicular to the membrane, and the cytoplasmic domain is wider than the extracellular domain, reminiscent of a truncated funnel as suggested by the 2D class averages (**Fig. 5h**). In the cytoplasmic domain, an open, dome-like pore entrance has a diameter of ∼42 Å (**Fig. 6b**). The walls of the dome are formed by the α-helices from the cytoplasmic loop and the C-terminus in the seven subunits. This pore diameter is similar to that observed for N-terminally truncated INX-6 (INX-6 ΔN) (Oshima et al., 2016a) but larger than that of wild-type INX-6 (INX-6 WT) (Oshima et al., 2016b) and the LRRC8A pore domain (Deneka et al., 2018) (**Table 1**). The narrowest constriction of the channel is at the entrance to the extracellular domain (**Fig. 6b**), similar to the constricted extracellular side of the LRRC8A pore domain (Deneka et al., 2018).

**Figure 6.**
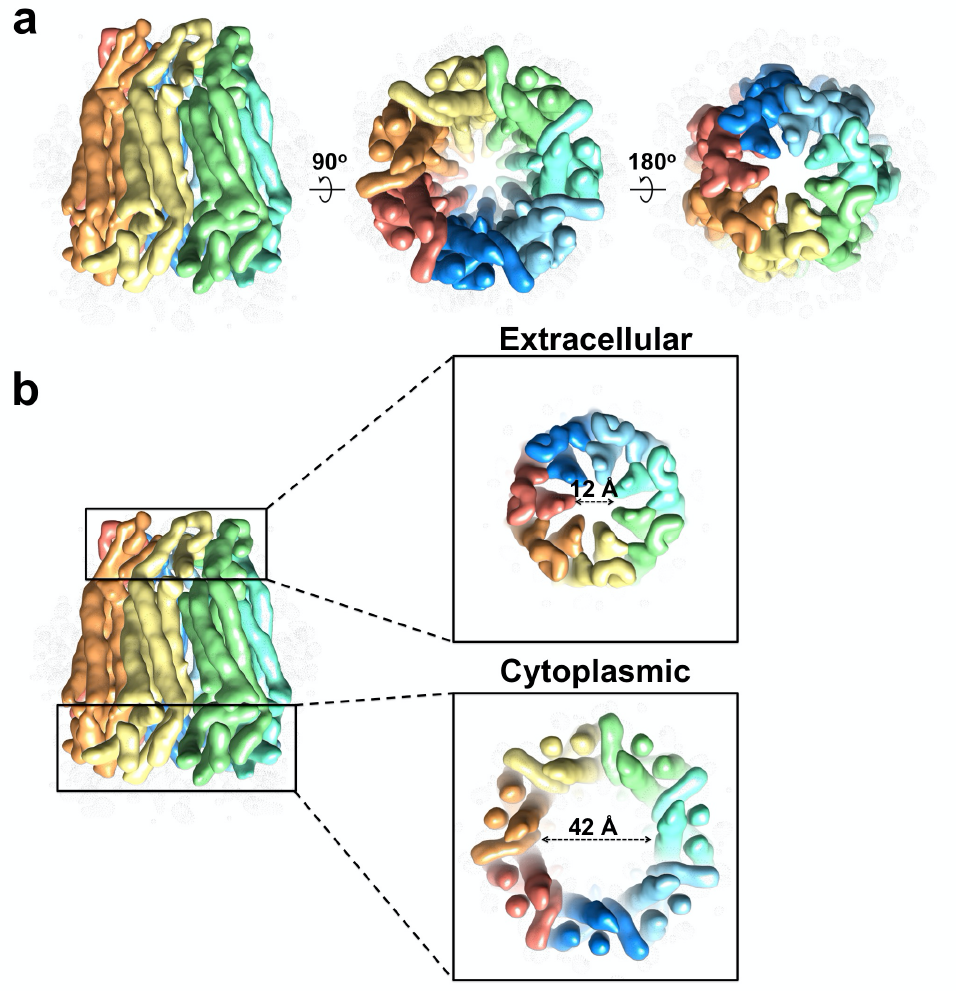
CryoEM density map of fPanx1ΔC in lipid nanodiscs displays a tripartite heptameric structure comprised of extracellular, TM and cytoplasmic domains. (**a**) Side and top (middle, from cytoplasmic surface; right, from extracellular surface) views of segmented cryoEM density map of fPanx1ΔC in lipid nanodiscs I which each of the 7 subunits is displayed in a different color. (**b**) Extracellular and cytoplasmic views of segmented cryoEM density map of fPanx1ΔC, with pore diameters of 12 Å and 42 Å, respectively.

## DISCUSSION

Over the last ∼15 years, Panx1 has been considered as a hexameric channel by analogy to Cx hemichannels (Baranova et al., 2004; Boassa et al., 2007; Bruzzone et al., 2003; Locovei et al., 2006a; Penuela et al., 2007), and based on previous empirical results from EM, chemical crosslinking, and single-molecule photobleaching (Ambrosi et al., 2010; Boassa et al., 2007; Chiu et al., 2017). In contrast, six publications in 2020 presented 16 independent, high-resolution cryoEM 3D reconstructions of various Panx1 constructs (Deng et al., 2020; Jin et al., 2020; Michalski et al., 2020; Mou et al., 2020; Qu et al., 2020; Ruan et al., 2020), which demonstrated that Panx1 assembles as a heptamer. In such studies the highest resolution is achieved by excluding the vast majority of the recorded particle images. Therefore, we performed physicochemical analysis to examine the entire population of particles and assess whether fPanx1ΔC may exist in multiple oligomeric states in solution. We also performed cryoEM in the membrane mimetics amphipol A8-35 and lipid bilayer nanodiscs. Use of a Volta phase plate enhanced the image contrast to facilitate particle picking. We examined a C-terminal truncation construct of frog Panx1 (fPanx1ΔC), which was basally closed and presumably exists in a single conformation.

### fPanx1ΔC in solution does not assemble as a hexamer

fPanx1ΔC formed thermostable and homogeneous oligomers when purified in a detergent mixture of n-dodecyl-β-D-maltopyranoside and cholesteryl hemisuccinate (DDM/CHS) (**Fig. 3a-c**). fPanx1ΔC was also homogeneous and monodisperse when reconstituted in the membrane mimetics amphipol A8-35 (**Fig. 4b,c,e**) and lipid nanodiscs (**Fig. 5b,d,g**). SDS-PAGE of fPanx1ΔC in DDM/CHS (**Fig. 3a**) and reconstituted in amphipol A8-35 **(Fig. 3a)** and nanodiscs (**Fig. 5a**) yielded an apparent monomer molecular weight of ∼35 kDa. Glutaraldehyde-crosslinked fPanx1ΔC in DDM/CHS (**Fig. 3a**) and reconstituted in lipid bilayer nanodiscs **(Fig. 5a)** migrated with an apparent molecular weight of ∼250 kDa, suggesting a heptameric oligomer (compared with ∼210 kDa for a hexamer). The electrophoretic mobility by SDS-PAGE (∼35 kDa) was smaller than the predicted mass of 43 kDa, presumably due to effects of the solubilizing agents. Therefore, we also performed SEC combined with multi-angle light scattering (SEC-MALS) (**Fig. 3f**). This enables an investigation of the stoichiometry of fPanx1ΔC by simultaneously measuring the protein and detergent components in a protein-detergent complex using a combination of SEC and light scattering with ultraviolet (UV) and refractive index (RI) detectors (Slotboom et al., 2008). SEC-MALS yielded an average mass of 320 kDa. Given the 5% experimental error of ∼16 kDa, the observed protein mass encompassed that predicted for a 301 kDa heptamer and a 344 kDa octamer, indicating that fPanx1ΔC *does not* assemble as a 258 kDa hexamer in solution in appreciable amounts.

### Negative-stain and cryoEM analysis demonstrated that fPanx1ΔC is heptameric in amphipol A8-35 and lipid nanodiscs

Frozen-hydrated fPanx1ΔC in DDM/CHS demonstrated low contrast in cryomicrographs due to high background noise contributed by free DDM/CHS micelles. To rectify this, fPanx1ΔC was exchanged into amphipol A8-35 as described previously (Althoff et al., 2011; Liao et al., 2013; Lu et al., 2014; Tribet et al., 1996). 2D class averages of negatively-stained and frozen-hydrated particles indicated a heptameric oligomer (**Fig. 4d,f**). However, an obstacle to 3D structure determination was that fPanx1ΔC in amphipols oriented preferentially in the vitreous ice of cryoEM grids, with the symmetry axis parallel to the incident electron beam (**Fig. 4f**). This obstacle was overcome by reconstituting fPanx1ΔC into lipid nanodiscs (**Figs. 5h and S3b**). Particle picking was greatly facilitated due to the high contrast provided by the VPP (**Fig. 5g**). The 3D reconstruction clearly showed that fPanx1ΔC is heptameric in which the TM protomer folds as a 4-helix bundle (**Figs. 5i,j,k and 6**). As is common with single-particle cryoEM, the 3D reconstruction was based on a small percentage (5.6%) of the particles (38,724 out of 687,949). Nevertheless, the 2D class averages did not suggest any other stoichiometries for the oligomer. Although the resolution of the cryoEM density map was limited to ∼7 Å, subnanometer resolution was sufficient to determine the heptameric stoichiometry of the channel and the 4-helix bundle design of the protomers (**Figs. 5k and 6**). In general, the oligomeric state, secondary and tertiary structure recapitulate that of the recent high-resolution cryoEM structures of Panx1 (Deng et al., 2020; Jin et al., 2020; Michalski et al., 2020; Mou et al., 2020; Qu et al., 2020; Ruan et al., 2020) (e.g., compare **Figs. 7a** and **7d**).

**Figure 7.**
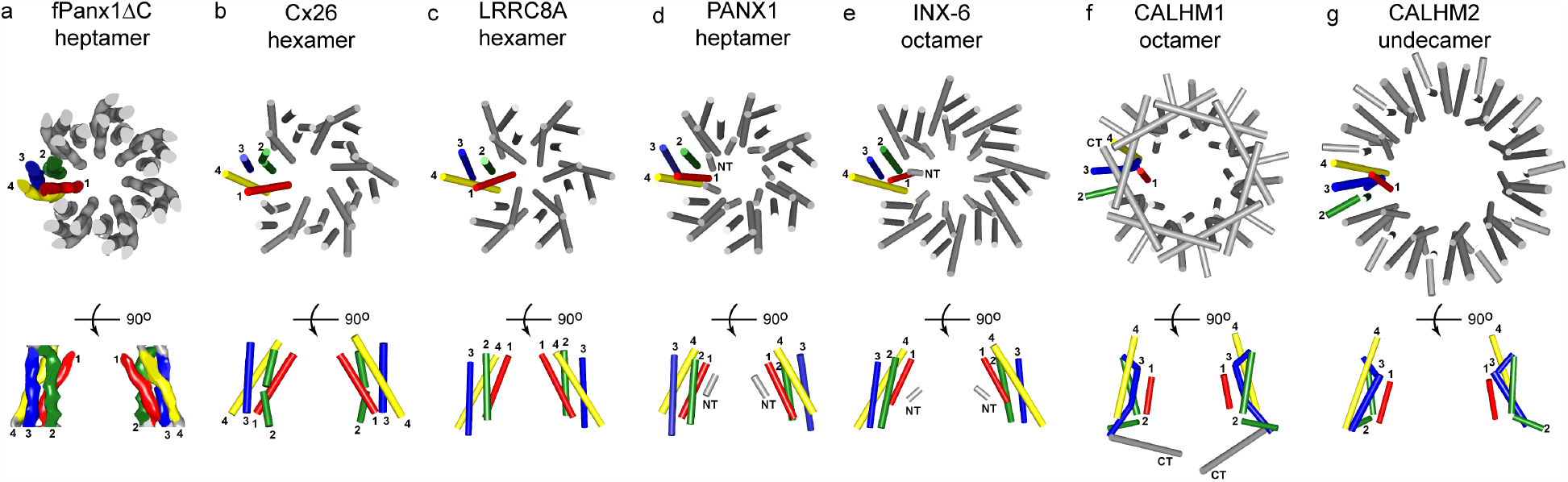
fPanx1ΔC has a similar transmembrane 4-helix bundle arrangement with Cx26, LRRC8A and INX-6, whereas CALHM1 and CALHM2 have a noncanonical 4-TM bundle. Pore diameter corresponds roughly with oligomeric state. Cross-sectional, oligomeric, top view (viewed from cytoplasmic side of membrane) and monomeric, side view of transmembrane (TM) α-helices from **(a)** Segmented cryoEM density map of fPanx1ΔC. (**b**)-(**g**) TM α-helices are shown as solid-cylinder representation. (**b**) hexamer of the Cx26 gap junction channel (PDB code: 5ER7) (Bennett et al., 2016), (**c**), hexameric LRRC8A (PDB code: 6g8z) (Deneka et al., 2018) (**d**) heptameric PANX1, (PDB code 6WBF) (Ruan et al., 2020), (**e**) octameric INX-6 (PDB code: 5h1q) (Oshima et al., 2016b) (**f**) octameric CALHM1 (PDB code: PDB: 6VAM) (Syrjanen et al., 2020a, b) and (**g**) (PBD code: 6uiv) (Choi et al., 2019).

### Comparison with other oligomeric, 4-helix bundle, large pore channels

Despite low sequence homology and different oligomeric states, the crossing angles of the Panx1 4-helix bundle are most similar to that of connexins, innexins and LRRC8 (SWELL) channels (**Fig. 7a-d**). The innermost pore lining helix is TM1, with TM2, TM3 and TM4 positioned in a counter-clockwise order when viewed from the cytoplasmic side of the membrane. By contrast, the TM helical arrangement in CALHM1 octamers and CALHM2 undecamers is clockwise, with TM2, TM3 and TM4 almost residing in a plane **(Fig. 7e,f)** (Choi et al., 2019). In spite of the variations in oligomeric state, all of the channels have a large cytoplasmic vestibule and narrow extracellular pore. The general shape of the domain architecture of the Panx1 protomer resembles that of the INX-6 hemichannel (Oshima et al., 2016b) and the LRRC8A channel pore domain (Deneka et al., 2018). The longitudinal height of fPanx1ΔC channel is ∼105 Å (**Fig. 5i,j**), which is also similar to the octameric INX-6 hemichannel (Oshima et al., 2016b) and the hexameric LRRC8A pore domain (Deneka et al., 2018) **(Table S4)**. In contrast, the cytoplasmic regions of Cx channels are largely disordered, which confounds precise measurement of the Cx channel dimensions (Bennett et al., 2016; Maeda et al., 2009; Unger et al., 1999). The diameter of fPanx1ΔC is greatest on the cytoplasmic surface (∼105 Å) (**Fig. 5i**), which is smaller than that of the octameric INX-6 (Oshima et al., 2016b) but larger than hexameric Cx hemichannels (Maeda et al., 2009) (**Table S4**), likely due to the different number and molar mass of protomers that comprise each channel.

The estimated diameter of the extracellular constriction (∼12 Å; **Fig. 6b**) is smaller than the effective diffusion diameter of ATP (∼14 Å). Considering the additional length contributed by amino-acid side chains, we infer that the extracellular pore is in a closed conformation, consistent with the basally closed state of the channel (**Fig. 2**). Interestingly, all of the high-resolution Panx1 structures display a narrow extracellular pore that would exclude ATP (Deng et al., 2020; Jin et al., 2020; Michalski et al., 2020; Qu et al., 2020; Ruan et al., 2020), even though constructs truncated at the C-tail caspase site (e.g., PDB 6WBG (Ruan et al., 2020) and PDB 6LTN (Mou et al., 2020)) should permit an open ATP-permeable conformation. Presumably, protein “breathing” would enable a wider extracellular pore that would accommodate hydrated ions, metabolites and signaling molecules like ATP.

The Panx1 heptamer provides further structural diversity within the family of large-pore channels, including hexameric Cx hemichannels, hexameric LRRC8 (SWELL1) channels, octameric Inx-6 hemichannels and CALHM1 channels, and undecameric CALHM2 channels. Conserved structural themes are a large cytoplasmic vestibule with a diameter that corresponds roughly with the oligomeric state and a 4-helix bundle protomer, albeit with noncanonical helical packing for CALHM1 and CALHM2.

## Acknowledgements

This work was supported by the National Institutes of Health (NIH) grants P01 HL120840 (D.A.B. and M.Y.), R01 HL48908 (M.Y.) and R01 GM138532. X.J. was supported by American Heart Association predoctoral fellowship 15PRE25560084. CryoEM data were collected at the University of Virginia Molecular Electron Microscopy Core, the National Cancer Institute’s National Cryo-Electron Microscopy Facility (NCEF) at the Frederick National Laboratory for Cancer Research (Sriram Subramaniam, Director), the Electron Bio-imaging Centre (eBIC) located at the Diamond Light Source (Peijun Zhang, Director), and the National Center for Macromolecular Imaging (NCMI) at Baylor College of Medicine (Wah Chiu, Director). We thank Maciej Jagielnicki for assistance with molecular graphics. Some graphics and analyses were performed with the University of California, San Francisco Chimera package. Chimera is developed by the Resource for Biocomputing, Visualization, and Informatics at the University of California, San Francisco (supported by the National Institute of General Medical Sciences Grant P41 GM103311).

## Author contributions

X.J. and S.A.L. performed protein expression and purification and amphipol and nanodisc reconstitution. B.C.B advised and technically supported protein purification. X.J. performed negative-stain EM and image analysis, cryoEM sample preparation and data acquisition, fluorescent thermal stability assay, protein crosslinking and the deglycosylation assay. X.J. and M.D.P. performed cryoEM and image analysis. W.E.M. performed MALDI-MS and SEC-MALS analysis. Y.-H.C. and D.A.B. performed electrophysiology. The manuscript was written primarily by X.J. and M.Y., with edits and contributions from all authors.

## Data and materials availability

The EM map of pannexin1 has been deposited in the EMDB (www.ebi.ac.uk/pdbe/emdb/) with accession code xxxx.

## METHOD DETAILS

### Mammalian cell culture and electrophysiology

HEK293T cells (ATCC, passage 3-14) were cultured at 37 °C with humidified air containing 5% CO_2_ in Dulbecco’s Modified Eagle Medium (DMEM, Gibco) containing 10% fetal bovine serum (FBS, Gibco), penicillin, streptomycin, and sodium pyruvate. Transfection was carried out using Lipofectamine2000™ (Invitrogen) according to the protocol provided by the manufacturer.

Whole-cell recordings were performed at room temperature using an Axopatch 200B amplifier and pCLAMP9 software (Molecular Devices). Ramp voltage commands were applied by using pCLAMP9 software and a Digidata 1322A digitizer (Molecular Devices). Borosilicate glass patch pipettes of 3-5 MΩ were formed using a micropipette puller (Sutter Instruments, P-97) and then coated with Sylgard 184 silicone (Dow Corning). HEK293T cells were plated onto poly-lysine-coated coverslips 16-18 h after transfection. The bath solution contained 10 mM HEPES, pH 7.3, 140 mM NaCl, 3 mM KCl, 2 mM MgCl_2_, 2 mM CaCl_2_ and 10 mM glucose. The pipette solution contained 10 mM HEPES, pH 7.3, 30 mM tetraethylammonium chloride, 100 mM cesium methanesulfonate, 4 mM NaCl, 1 mM MgCl_2_, 0.5 mM CaCl_2_, 10 mM EGTA, 3 mM ATP-Mg and 0.3 mM GTP-Tris. Carbenoxolone (CBX, 50 µM) was applied in bath solutions after acquisition of steady-state fPanx1ΔC activities. Phenylephrine (PE, 20 µM in the bath solution) was used to induce α1D adrenergic receptor-mediated activation of fPanx1ΔC channels (Billaud et al., 2015; Billaud et al., 2011; Chiu et al., 2017). Data were filtered at 1 kHz and digitized at a sampling rate of 2 kHz. Data were analyzed using pCLAMP and GraphPad Prism software, and the results are presented as mean ± S.E.M.

### Protein expression and purification

Beginning in 2009, we examined 22 constructs of human, mouse and frog Panx1 (**Table S1**). Protein expression and purification were performed as previously described (Chiu et al., 2017), with modifications. The construct that displayed a high level of expression and remained homogenous and monodisperse during purification was a C-terminal truncation (deletion from residue 358 to 428) of frog Panx1 (NCBI: NP_001123728, 428 amino acids), modified with a thrombin cleavage recognition sequence (LVPRGS) followed by a strep tag II (WSHPQFEK) at the carboxy-terminus (fPanx1ΔC). The construct was subcloned into the pFastBac1 vector (Invitrogen) for baculovirus expression in *Spodoptera frugiperda* (Sf9) insect cells using the Bac-to-Bac expression system (Invitrogen). Recombinant fPanx1ΔC baculovirus was used to infect Sf9 insect cells grown at 27 °C to a density of 2 × 10^6^ ml^-1^, at a multiplicity of infection (MOI) of ∼3. Cells were collected 48 h after infection by low-speed centrifugation at 2,000 x g. To isolate membrane-localized fPanx1ΔC, Sf9 cell pellets were resuspended in low salt buffer (50 mM HEPES, pH 7.5, 50 mM NaCl, 0.5 mM EDTA and protease inhibitor cocktails (Roche) and lysed by Dounce homogenization (∼30 strokes). Nucleic acids were digested by adding MgCl_2_ to 2.5 mM and Benzonase (EMD Millipore) to ∼12.5 units per 1 ml lysate, with gentle stirring at 4 °C for 20 min. Membranes were collected by ultracentrifugation at 100,000 x g and washed with stepwise Dounce homogenization in low salt buffer and high salt buffer (50 mM HEPES, pH 7.5, 1 M NaCl, 0.5 mM EDTA and protease inhibitor cocktails). Pellets were isolated by ultracentrifugation at 100,000 x g between steps. The final Dounce homogenization was performed in intermediate salt buffer (50 mM HEPES, pH 7.5, 500 mM NaCl), followed by ultracentrifugation at 100,000 x g. The membrane pellet was solubilized at 4 °C for 4 h with 1% (w/v) n-dodecyl-β-D-maltopyranoside (DDM, Anatrace) and 0.2% (w/v) cholesteryl hemisuccinate (CHS, Anatrace) in ∼50 ml of buffer containing 50 mM HEPES, pH 7.5, 300 mM NaCl, 3 mM CaCl_2_, 2.5% glycerol and protease inhibitor cocktails. Insoluble material was removed by ultracentrifugation at 100,000 x g, and the supernatant was incubated with ∼1.0 ml of Strep-Tactin Superflow Plus resin (QIAGEN) overnight at 4 °C. The resin was packed in an Econo-column (Bio-Rad, 1.0 x 10 cm) and washed with high salt buffer (50 mM HEPES, pH 7.5, 1 M NaCl, 3 mM CaCl_2_ and 0.05% DDM with 0.01% CHS), for 20 column volumes/wash, and eluted with 2.5 mM Desthiobiotin (Sigma) in buffer (50 mM HEPES, pH 7.5, 500 mM NaCl, 3 mM CaCl_2_ and 0.02% DDM with 0.004% CHS). The eluted protein was concentrated to 2-3 mg ml^-1^ using an Amicon ultracel-100 centrifugal filter unit (EMD Millipore) for amphipol exchange or nanodisc reconstitution. Analytical size-exclusion chromatography (SEC) was performed on a Superpose 6 increase 10/300 GL column (GE Healthcare) interfaced to an AKTA Purifier10 HPLC system (GE Healthcare), equilibrated with buffer (50 mM HEPES, pH 7.5, 500 mM NaCl, 3 mM CaCl_2_ and 0.02% DDM with 0.004% CHS).

### Amphipol exchange and nanodisc reconstitution

For amphipol exchange, purified fPanx1ΔC in DDM/CHS was mixed with amphipathic surfactant Amphipol A8-35 (Anatrace) at a ratio of 1:3 (*w/w*) by gentle agitation for 4 h at 4 °C. The DDM/CHS detergent was then removed by overnight incubation with Bio-Beads SM-2 (15 mg per 1 ml protein/detergent/amphipols mixture, Bio-Rad) at 4 °C, and the Bio-Beads were subsequently removed using a disposable polyprep column (Bio-Rad).

For nanodisc reconstitution, membrane scaffold protein 2N2 (MSP2N2) was expressed and purified from *Escherichia coli* as previously described (Ritchie et al., 2009), with modifications. The purified MSP2N2 was solubilized in buffer (50 mM Tris pH 8.0, 100 mM NaCl, 4 mM β-mercaptoethanol, 2 mM EGTA and 10 mM sodium cholate). 2.5 mg soybean polar lipid extract (Avanti) dissolved in chloroform was dried under an argon stream to form a lipid film. Residual chloroform was further removed by vacuum desiccation for 3 h. The lipid film was then rehydrated in buffer (25 mM Tris pH 7.5, 150 mM NaCl, 2% n-octyl-β-D-glucopyranoside (OG) (*w/v*)) by vigorous mixing, resulting in a clear lipid stock at 10 mM concentration. Purified fPanx1ΔC (2-3 mg ml^-1^, 45-70 μM) solubilized in DDM/CHS was mixed with the soybean lipid stock (7.5 mg ml^-1^, 10 mM) and MSP2N2 (2.4 mg ml^-1^, 56 μM) at a molar ratio of 1:1.25:125 (fPanx1ΔC monomer:MSP2N2:soybean lipid) and incubated on ice for 1 h. Bio-beads SM2 (10 mg ml^-1^, Bio-Rad) were added to initiate the reconstitution by removing detergents from the system, and the mixture was incubated at 4 °C for 1 h with constant rotation. A second aliquot of Bio-beads was then added to increase the concentration to 30 mg ml^-1^, and the sample was incubated at 4 °C overnight with constant rotation. Bio-beads were subsequently removed using a disposable polyprep column (Bio-Rad).

fPanx1ΔC in amphipols or nanodiscs was further purified by size-exclusion chromatography (SEC). Insoluble material was removed by ultracentrifugation at 150,000 x g for 20 min before purification by SEC on a Superpose 6 increase 10/300 GL column (GE Healthcare) equilibrated with buffer (25 mM HEPES, pH 7.5, 150 mM NaCl and 3 mM CaCl_2_). The peak fractions containing fPanx1ΔC were collected and concentrated to ∼0.2 mg ml^-1^ using a 0.5 ml concentrator with 100 kDa cut-off (EMD Millipore), for negative-stain EM and cryoEM. In addition to SEC, fPanx1ΔC purity was assessed by SDS–polyacrylamide gel electrophoresis (PAGE) using 4-20% pre-cast gradient Tris-glycine gel (Bio-Rad) stained with Simply Blue (SimpleBlue Safe Stain, Novex), and immunoblotting using anti-Strep antibodies (Qiagen).

### Fluorescent thermal stability assay (FTSA)

The thermal stability of fPanx1ΔC in the detergent DDM/CHS was characterized using a cysteine-reactive, coumarin-based fluorophore, CPM (N-(4-(7-diethylamino-4-methyl-3-coumarinyl) phenyl) maleimide (Alexandrov et al., 2008). The quantum yield increases upon temperature-induced protein unfolding when CPM binds to cysteine residues within the hydrophobic region of a protein. The fPanx1ΔC contains 8 cysteine residues: 4 Cys in the extracellular loops (EL) (C66 and C84 in EL1; C248 and C267 in EL2), 3 Cys in the TM domains (C40 in TM1, C218 and C230 in TM3) and 1 Cys in the cytoplasmic loop (CL) (C149). The 3 Cys residues in the TM domains are predicted to be free cysteines that are maintained in a reduced state, which would be potentially accessible for binding to CPM as the protein is thermally denatured. When bound to a Cys thiol, the emission wavelength is 463 nm.

The CPM fluorescent dye (Invitrogen) dissolved in dimethylformamide was diluted with SEC buffer (50 mM HEPES, pH 7.5, 500 mM NaCl, 3 mM CaCl_2_ and 0.02% DDM with 0.004% CHS) to 13.3 μM and incubated with buffer on ice for 15 min. An aliquot (10 μg) of fPanx1ΔC was added to the buffer containing CPM fluorescent dye and incubated on ice for another 15 min. The temperature scan from 10 to 90 °C was performed using a FluoroMax-3 spectrofluorometer (Horiba Jobin-Yvon), with excitation and emission wavelengths of 387 nm and 463 nm, respectively. The fluorescence-temperature profile was analyzed using non-linear regression of a Boltzmann sigmoidal equation (Origin 7.5 software, OriginLab). The melting temperature (T_m_) was calculated from the inflection point of the resulting melting curve as described previously (Alexandrov et al., 2008).

### Protein crosslinking

Purified fPanx1ΔC (1-2 mg ml^-1^ in buffer composed of 50 mM HEPES pH 7.5, 500 mM NaCl, 3 mM CaCl_2_, 0.02% DDM with 0.004% CHS) was crosslinked by 0.1% glutaraldehyde, a nonspecific crosslinker of amine-to-amine lysine residues. After 15 min incubation at room temperature, the reaction was terminated by addition of 100 mM Tris-HCl, pH 8.0. The crosslinking was confirmed by Coomassie blue stained SDS-PAGE and Westerm immunoblot analysis using anti-Strep antibodies.

### Deglycosylation assay

Purified fPanx1ΔC (2-3 mg ml^-1^ in buffer composed of 50 mM HEPES, pH 7.5, 500 mM NaCl, 3 mM CaCl_2_ and 0.02% DDM) was incubated with 2000 units of N-glycosidase F (PNGase F) (New England BioLabs) at room temperature for 1 h. PNGase F was removed by SEC on a Superpose 6 increase 10/300 GL column (GE Healthcare) equilibrated with buffer (50 mM HEPES, pH 7.5, 500 mM NaCl, 3 mM CaCl_2_ and 0.02% DDM). Deglycosylation was confirmed by SDS-PAGE, and the SEC peak fractions corresponding to deglycosylated fPanx1ΔC oligomer were collected and concentrated to ∼1 mg ml^-1^ using a 0.5 ml concentrator with 100 kDa cut-off (EMD Millipore). Such samples were used for MALDI-MS and SEC-MALS.

### Matrix-assisted laser desorption/ionization mass spectrometry (MALDI-MS)

Purified fPanx1ΔC (∼20 μM in buffer composed of 50 mM HEPES, pH 7.5, 500 mM NaCl, 3 mM CaCl_2_ and 0.02% DDM) was mixed with an aromatic carboxylic acid matrix, spotted onto the matrix-assisted laser desorption/ionization (MALDI) sample plate, and then loaded into the mass spectrometer. MALDI-MS analysis was performed by the Biomolecular Analysis Facility Core at the University of Virginia.

### Size-exclusion chromatography with multi-angle light scattering (SEC-MALS)

SEC-MALS experiments were performed at room temperature using an HPLC system, equipped with a UV detector, a miniDAWN TREOS MALS detector and an Optilab T-rEX refractive index detector (Wyatt Technologies). An aliquot (∼100 μg) of purified, deglycosylated fPanx1ΔC in buffer (50 mM HEPES, pH 7.5, 500 mM NaCl, 3 mM CaCl_2_ and 0.02% DDM) was injected onto a Superdex 200 Increase 10/300 GL column (GE Healthcare), equilibrated with buffer (25 mM HEPES, pH 7.5, 150 mM NaCl, 3 mM CaCl_2_ and 0.02% DDM). The data were analyzed using Astra software (Astra 6.1, Wyatt Technologies) on the basis of the absorption at 280 nm, light scattering, and the differential refractive index. Extinction coefficients at 280 nm and *dn/dc* values for Panx1 were estimated to be 1.472 ml mg^-1^ cm^-1^ and 0.185 mol ml g^-2^, respectively. The *dn/dc* value for DDM was assumed to be 0.133 mol ml g^-2^ (Strop and Brunger, 2005). The system was normalized using Bovine Serum Albumin (BSA, Sigma).

### Negative-stain EM and image analysis

An aliquot (3.5 μl) of purified fPanx1ΔC complex (0.01-0.02 mg ml^-1^) was applied to a glow-discharged, copper grid covered with a thin layer of continuous carbon film (300-mesh, Electron Microscopy Sciences), and stained with 2% (w/v) uranyl acetate (Adair and Yeager, 2007). Negatively-stained EM grids were imaged on a Tecnai F20 electron microscope (FEI Company), operating at 120 kV. Images were recorded at a nominal magnification of ×62,000 and a defocus of −0.75 μm using a 4K × 4K charge-coupled device (CCD) camera (UltraScan 4000, Gatan), corresponding to a calibrated pixel size of 1.82 Å on the specimen.

EMAN2 software (Ludtke et al., 1999) was used for single-particle image analysis. To improve the signal-to-noise ratio and facilitate particle picking, negatively-stained electron micrographs were high-pass (100 Å) and low-pass (10 Å) Gaussian-filtered. A total of 11,275 fPanx1ΔC-amphipol particles from 55 micrographs, 14,144 fPanx1ΔC-nanodisc particles from 108 micrographs, and 8,135 crosslinked fPanx1ΔC-nanodisc particles from 110 micrographs, were semi-automatically selected using the ‘Swarm’ tool in the e2boxer.py program of EMAN2, and extracted within boxes of 196 pixels × 196 pixels. The contrast transfer function (CTF) was estimated and corrected by the e2ctf.py program of EMAN2. Particle images were normalized, centered and subjected to 8 cycles of 2D classification using the e2refine2d.py program by iterative, multivariate statistical analysis (MSA). Data collection statistics for negative-stain EM are summarized in **Table S2**.

### CryoEM sample preparation and data acquisition

An aliquot (3 μl) of purified fPanx1ΔC in amphipol A8-34 or lipid nanodiscs (∼0.2 mg ml^-1^ in buffer composed of 25 mM HEPES, pH 7.5, 150 mM NaCl and 3 mM CaCl_2_) was applied to a glow-discharged (with amylamine) C-flat holey carbon grid (1.2 μm hole size and 1.3 μm hole spacing on 400-mesh copper grid). The grid was blotted with Whatman #1 filter paper for 7.5 s using a Vitrobot Mark IV (FEI) maintaining at 100% humidity and 4 °C with a blot force of 2, and the grid was plunge-frozen into liquid ethane cooled by liquid nitrogen.

CryoEM was performed at the University of Virginia Molecular Electron Microscopy Core, the National Cancer Institute’s National Cryo-Electron Microscopy Facility (NCEF) at the Frederick National Laboratory for Cancer Research, the Electron Bio-imaging Centre (eBIC) located at the Diamond Light Source, and the National Center for Macromolecular Imaging (NCMI) at Baylor College of Medicine. Grids containing fPanx1ΔC-nanodisc complexes with optimal ice thickness were imaged at the National Cancer Institute’s National Cryo-Electron Microscopy Facility (NCI-NCEF) at the Frederick National Laboratory for Cancer Research using a Titan Krios electron microscope (Thermo Fisher Scientific), operated at 300 kV, equipped with a K2 Summit direct electron detector (Gatan), a GIF quantum energy filter (Gatan) with a 20 eV slit and zero-energy-loss mode to remove inelastic scattering, and a Volta phase plate (Thermo Fisher Scientific). A total of 3,566 cryomicrographs were recorded using an automated acquisition program EPU (Thermo Fisher Scientific), in EFTEM nanoprobe mode, using a 70 μm C2 aperture, at a nominal magnification of x105,000, corresponding to a calibrated physical pixel size of 1.32 Å per pixel on the specimen. Images were recorded at a defocus of −0.5 μm in counting mode, in which each image was fractionated into 40 frames with a total exposure time of 10 s. At a dose rate of 6.97 e^-^/pixel/sec, the total accumulated dose was 40 e^-^/Å^2^.

### CryoEM data processing

All image processing was performed using RELION 2.1 (Scheres, 2012). Movie frames (3-40) were used for correction of beam-induced motion and dose-weighting using MotionCor2 (Zheng et al., 2017), with 5 x 5 patches and the corresponding dose per frame. The contrast transfer function (CTF) parameters and phase shift were estimated using Gctf-v1.18 (Zhang, 2016). Low-quality micrographs with considerable drift, high defocus or poor fitting of the CTF were discarded, resulting in a dataset of 2,644 images for further image processing. 2,199 particles were manually picked and subjected to an initial reference-free 2D classification (10 classes), in which five distinctive 2D class averages were selected and used as templates for automated particle picking in RELION (Scheres, 2015). Auto-picked particles on each micrograph were manually screened to discard wrongly picked ice contamination, extracted with a box size of 180 x 180 pixels, and sorted by similarity to reference images to discard those with low z-scores. A starting dataset of 687,949 particles was generated for 2D classification. After one round of 2D classification (200 classes), false positive and poor quality particles were discarded, resulting in 319,685 particles for 3D classification. CryoSPARC (Punjani et al., 2017) was used to analyze a subset of 84,415 particles from 2D classification in order to generate an *ab initio* model. RELION 2.1 (Scheres, 2012) was then used to perform 3D classification (1 class, C7 symmetry) and manual refinement by adjusting the angular sampling degree (from 7.5° to 3.7° to 1.8°). The initial model was then low-pass filtered to 50 Å in RELION 2.1 (Scheres, 2012), which was used as a reference in subsequent 3D classification. The heptameric structure was evident from 2D classes. We therefore performed one round of 3D classification (10 classes), imposing C7 symmetry to identify a homogenous subset of particles. Two out of ten 3D classes showed densities consistent with TM α-helices. Class 6 displayed a larger diameter nanodisc compared to Class 8. An angular distribution plot showed that the 3D class of the larger diameter nanodiscs contained primarily top views, whereas the 3D class of the smaller diameter nanodiscs displayed an even particle distribution of both top and side views. Thus, 46,450 particles in the small-nanodisc 3D class were subjected to another round of 2D classification (50 classes), resulting in 38,724 particles for 3D auto-refinement, applying C7 symmetry. The resolution of the auto-refined 3D map before post-processing (unmask) was 7.7 Å on the basis of the 0.143 gold-standard Fourier Shell Correlation (FSC). RELION post-processing, with application of a soft mask, improved the resolution of the final map to 7.0 Å, and the map was sharpened with a B-factor of −899 Å^2^. Statistics for cryoEM data collection and image processing are summarized in **Table S3**. UCSF Chimera (Pettersen et al., 2004) was used to visualize the cryoEM density map, which was segmented using the ‘Segment map’ function.

## SUPPLEMENTAL FIGURES AND TABLES

**Figure S1.**
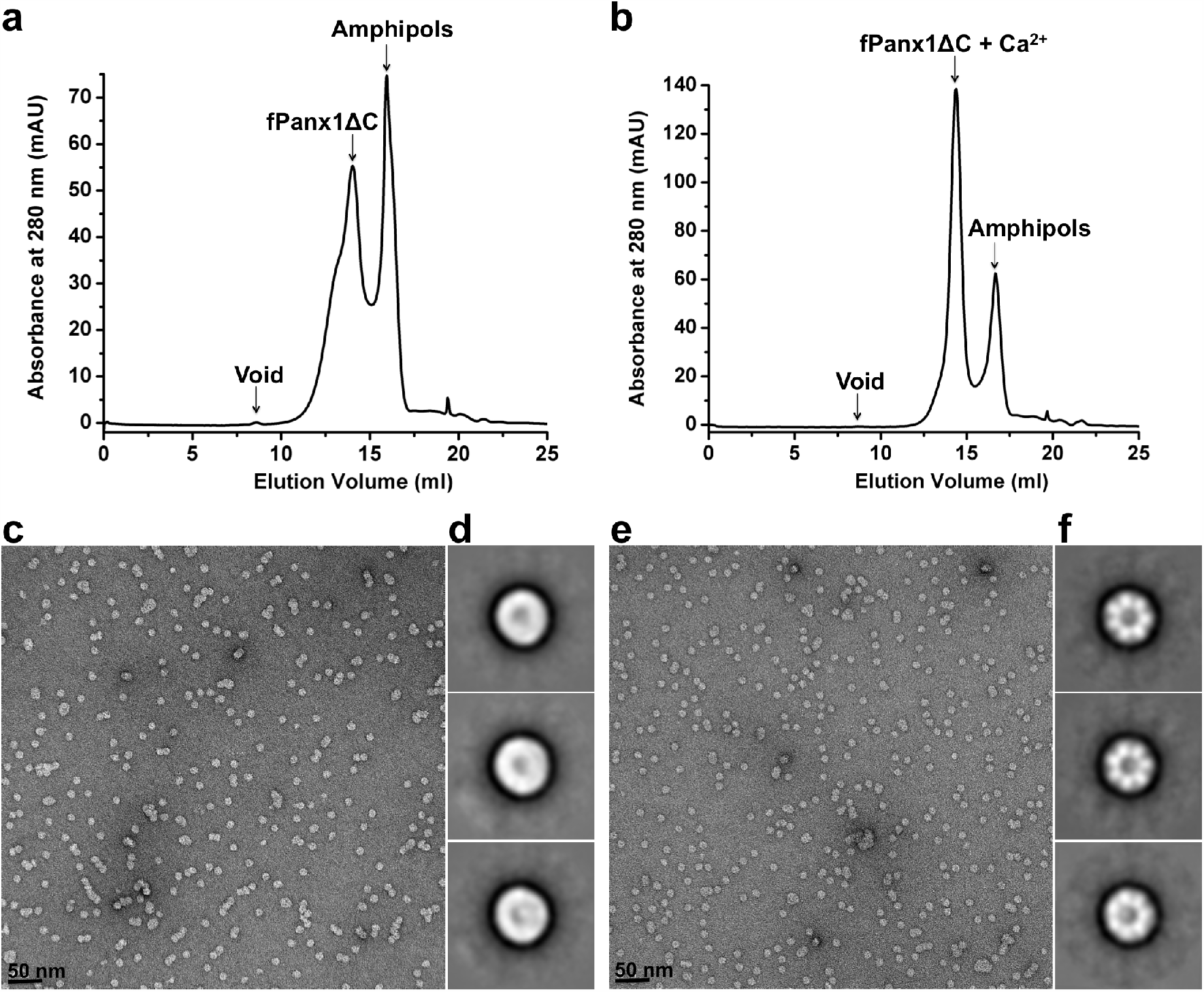
Calcium improved the homogeneity and heptameric symmetry of fPanx1ΔC particles, as characterized by SEC and single-particle analysis of negatively-stained particles. SEC of fPanx1ΔC in the absence **(a)** and presence **(b)** of Ca^2+^. The fPanx1ΔC peak was more homogenous in the presence of Ca^2+^ and lacked the left-sided ‘shoulder’ seen in absence of Ca^2+^. fPanx1ΔC protein, amphipols and void peaks are indicated. (**c,e**) Electron micrographs of negatively-stained fPanx1ΔC without Ca^2+^ (**c**) and with Ca^2+^ (**e**). Scale bar, 50 nm. (**d,f**) Representative 2D class averages of negatively-stained fPanx1ΔC without Ca^2+^ (**d**) and with Ca^2+^ (**f**). The particle box dimension is 357 Å.18,623

**Figure S2.**
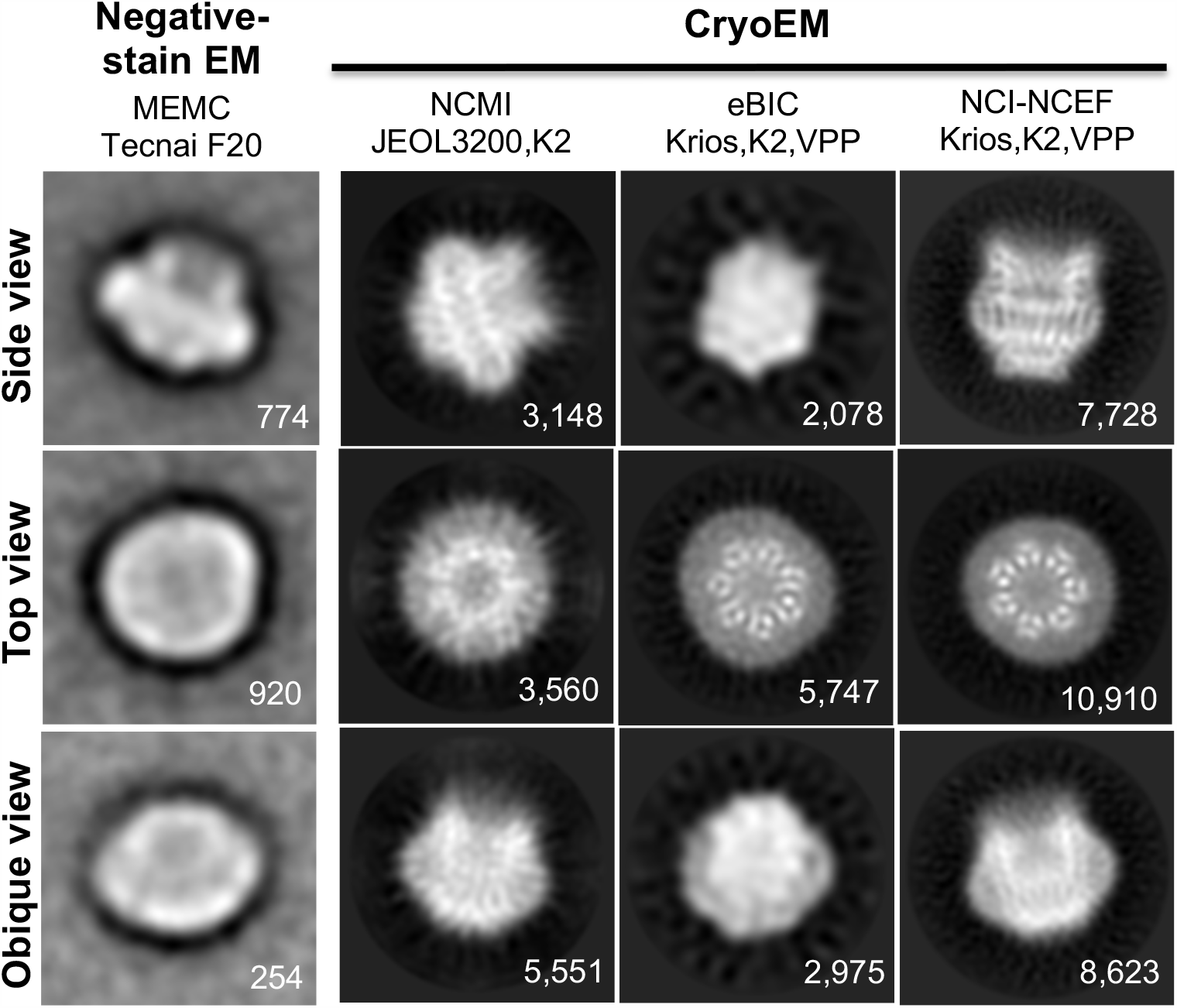
Progress on single-particle cryoEM of fPanx1ΔC in lipid bilayer nanodiscs. Negative-stain EM of fPanx1ΔC-nanodisc displayed characteristic channel features and different particle orientations, indicating that the sample was suitable for 3D reconstruction by single-particle cryoEM. We first collected cryomicrographs of Panx1 in nanodiscs at NCMI and eBIC using a K2 direct electron detector. Both data sets displayed circular densities in top view class averages consistent with end-on views of α-helices. Importantly, by using the phase plate, the top view class averages derived from the eBIC data showed better definition of α-helical secondary structure compared with the NCMI data, in which the heptameric arrangement of channel subunits with α-helices were clearly observed; however, the side view of the eBIC data did not reveal any secondary structural details, presumably due to the vitreous ice being too thick. By using a grid with optimally thin ice, we collected a set at NCI-NCEF using a Titan Krios equipped with a Gatan K2 Summit direct electron detector and a Volta phase plate. 2D class averages derived from the NCEF data set showed well delineated α-helices in both top and side views.

**Figure S3.**
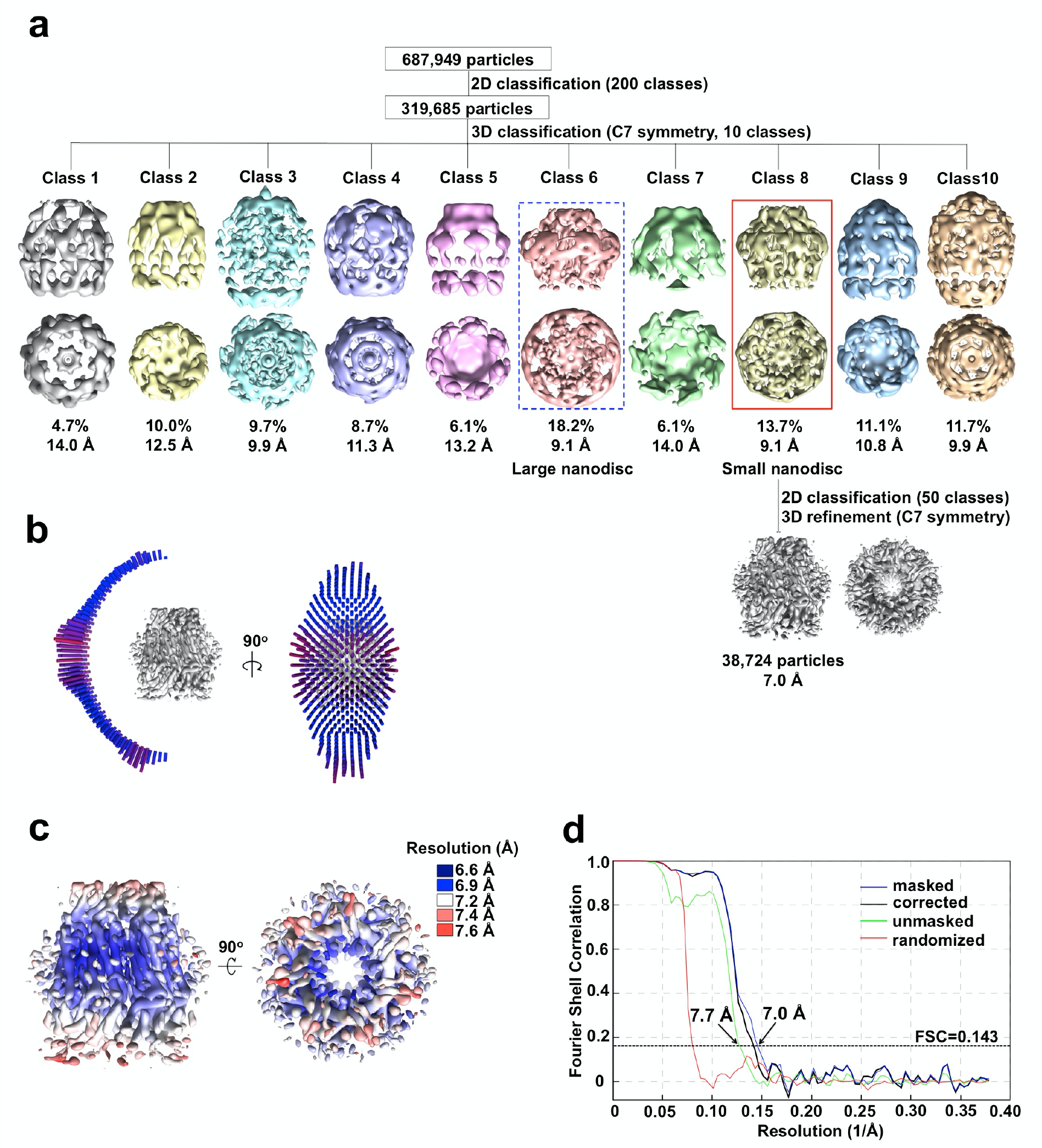
Single-particle cryoEM analysis of fPanx1ΔC in lipid nanodiscs. (**a**) Image processing workflow. With applying C7 symmetry, two (Class 6 and Class 8) out of ten classes generated from 3D classification revealed α-helical structures in the transmembrane domain, with two sizes of nanodiscs: a large nanodisc (Class 6, blue box) and a small nanodisc (Class 8, red box). Particles assigned to the small-nanodisc class (Class 8) were used for further refinement. The distribution of all particles (%) and the resolution of each class are indicated. (**b**) Angular distribution plot of all particles included in the final C7-symmetrized 3D reconstruction, showing both side and top view orientations. The length and the color of cylinders correspond to the number of particles with respective Euler angles. **(c)** Final 3D reconstruction was colored according to the estimation of local resolution, shown as side and top views. **(d)** Fourier shell correlation (FSC) curves of the final refined masked (blue), unmasked (green), phase-randomized (red) and corrected for mask convolution effects (black) cryoEM density map.

**Table S1.**
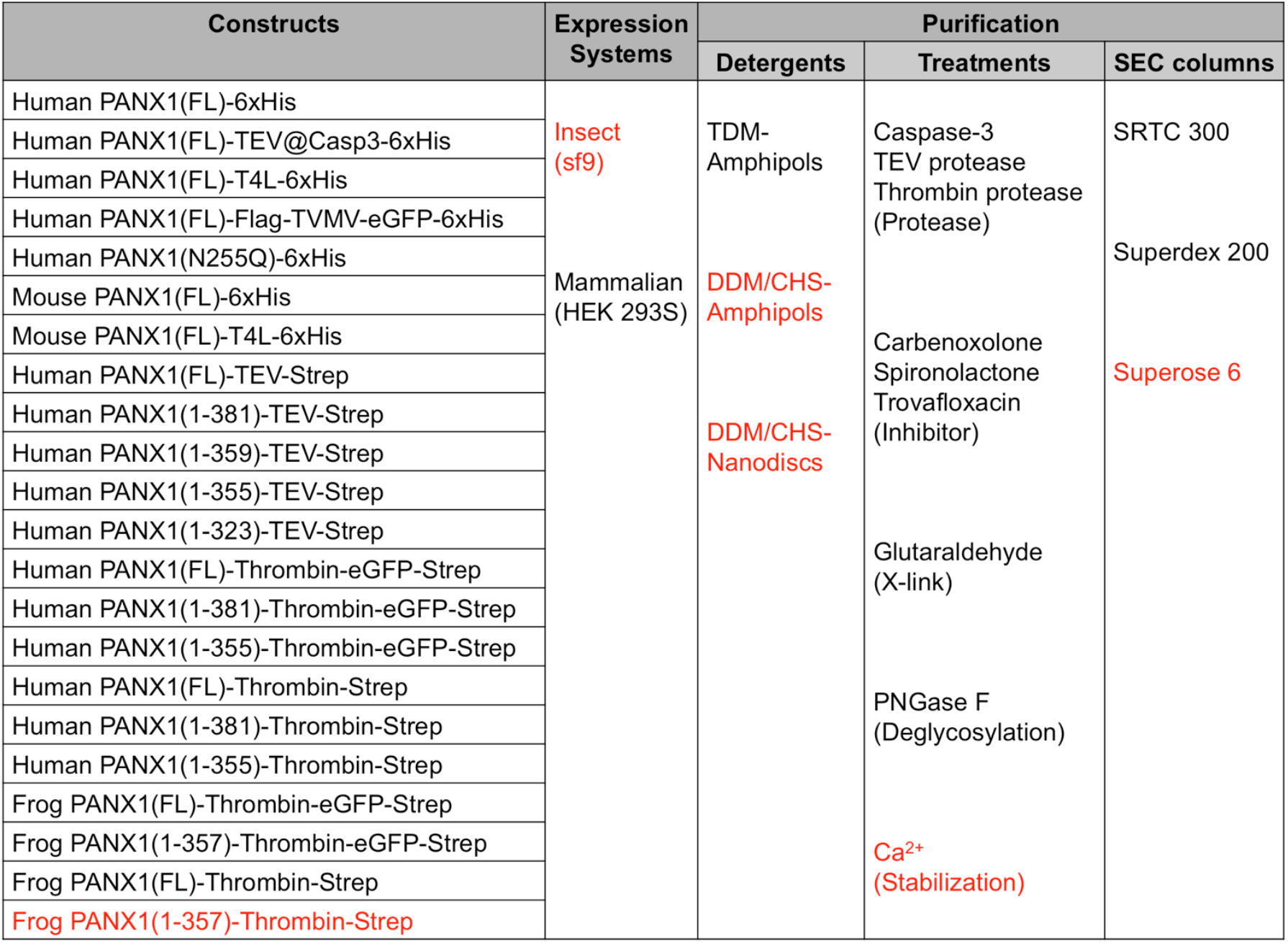
Optimization of Panx1 Samples for CryoEM and Single-Particle Image Analysis. Final conditions are highlighted in red.

**Table S2.** Summary of data collection statistics for negative-stain EM analysis.

| **Negative-stain EM**<br>**data collection and processing** | **fPanx1ΔC**<br>**in amphipols** | **fPanx1ΔC**<br>**in nanodiscs** | **X-linked fPanx1ΔC**<br>**in nanodiscs** |
| --- | --- | --- | --- |
| Electron microscope | Tecnai F20 | Tecnai F20 | Tecnai F20 |
| Electron detector | Gatan CCD (4k x 4k) | Gatan CCD (4k x 4k) | Gatan CCD (4k x 4k) |
| Voltage (kV) | 120 | 120 | 120 |
| Magnification | 62,000 | 62,000 | 62,000 |
| Defocus range (μm) | -0.75 | -0.75 | -0.75 |
| Pixel size (Å) | 1.82 | 1.82 | 1.82 |
| Total electron dose (e<sup>-</sup>/Å<sup>2</sup>) | 15 | 15 | 15 |
| Exposure time (s) | 1 | 1 | 1 |
| Number of images | 55 | 108 | 110 |
| Number of particles | 11,275 | 14,144 | 8,135 |
| Number of classes | 8 | 16 | 12 |
| Box size (pix) | 196 x 196 | 196 x 196 | 196 x 196 |

**Table S3.** Summary of data collection and statistics by single-particle cryoEM analysis.

| **CryoEM**<br>**data collection and processing** | **fPanx1ΔC** |
| --- | --- |
| Electron microscope | Titan Krios |
| Electron detector | K2 Summit |
| Voltage (kV) | 300 |
| Magnification | 105,000 |
| Defocus range (μm) | -0.3 to -1.2 |
| Pixel size (Å) | 1.32 |
| Total electron dose (e<sup>-</sup>/Å<sup>2</sup>) | 40 |
| Exposure time (s) | 10 |
| Number of images | 3566 |
| Number of frames per image | 40 |
| Phase plate periodicity | 25 |
| Initial particle number | 687,949 |
| Final particle number | 38,724 |
| Symmetry imposed | C7 |
| Map resolution (Å) | 7.0 |
| Map sharpening B factor (Å<sup>2</sup>) | -898 |

**Table S4.** Comparison of channel dimensions and pore diameters of fPanx1ΔC with other 4-helix bundle, large pore channels.

| Structures | Cytoplasmic diameter | (Hemi)channel height | Cytoplasmic pore diameter | Extracellular pore diameter |
| --- | --- | --- | --- | --- |
| Cx43(263T) (Unger *et al.*, 1999) | 70 Å | 75 Å | 40 Å | 15 Å |
| Cx26 (Maeda *et al.*, 2009) | 92 Å | 80 Å | 40 Å | 14 Å |
| INX-6ΔN (Oshima *et al.*, 2016) | 115 Å | 120 Å | 40 Å | 30 Å |
| INX-6WT (Oshima *et al.*, 2016) | 110 Å | 100 Å | 30 Å | 19 Å |
| LRRC8A (PD) (Deneka *et al.*, 2018) | 110 Å | 100 Å | 32 Å | 6 Å |
| fPannx1ΔC | 105 Å | 105 Å | 42 Å | 12 Å |

## REFERENCES

Abascal, F., and Zardoya, R. (2012). LRRC8 proteins share a common ancestor with pannexins, and may form hexameric channels involved in cell-cell communication. Bioessays 34, 551–560.

Adair, B.D., and Yeager, M. (2007). Electron microscopy of integrins. Methods Enzymol 426, 337–373.

Adamson, S.E., Meher, A.K., Chiu, Y.H., Sandilos, J.K., Oberholtzer, N.P., Walker, N.N., Hargett, S.R., Seaman, S.A., Peirce-Cottler, S.M., Isakson, B.E., et al. (2015). Pannexin 1 is required for full activation of insulin-stimulated glucose uptake in adipocytes. Mol Metab 4, 610–618.

Alexandrov, A.I., Mileni, M., Chien, E.Y., Hanson, M.A., and Stevens, R.C. (2008). Microscale fluorescent thermal stability assay for membrane proteins. Structure 16, 351–359.

Althoff, T., Mills, D.J., Popot, J.L., and Kühlbrandt, W. (2011). Arrangement of electron transport chain components in bovine mitochondrial supercomplex I1III2IV1. EMBO J 30, 4652–4664.

Ambrosi, C., Gassmann, O., Pranskevich, J.N., Boassa, D., Smock, A., Wang, J., Dahl, G., Steinem, C., and Sosinsky, G.E. (2010). Pannexin1 and Pannexin2 channels show quaternary similarities to connexons and different oligomerization numbers from each other. J Biol Chem 285, 24420–24431.

Bao, L., Locovei, S., and Dahl, G. (2004). Pannexin membrane channels are mechanosensitive conduits for ATP. FEBS Lett 572, 65–68.

Baranova, A., Ivanov, D., Petrash, N., Pestova, A., Skoblov, M., Kelmanson, I., Shagin, D., Nazarenko, S., Geraymovych, E., Litvin, O., et al. (2004). The mammalian pannexin family is homologous to the invertebrate innexin gap junction proteins. Genomics 83, 706–716.

Bennett, B.C., Purdy, M.D., Baker, K.A., Acharya, C., McIntire, W.E., Stevens, R.C., Zhang, Q., Harris, A.L., Abagyan, R., and Yeager, M. (2016). An electrostatic mechanism for Ca(2+)-mediated regulation of gap junction channels. Nat Commun 7, 8770.

Billaud, M., Chiu, Y.H., Lohman, A.W., Parpaite, T., Butcher, J.T., Mutchler, S.M., DeLalio, L.J., Artamonov, M.V., Sandilos, J.K., Best, A.K., et al. (2015). A molecular signature in the pannexin1 intracellular loop confers channel activation by the α1 adrenoreceptor in smooth muscle cells. Sci Signal 8, ra17.

Billaud, M., Lohman, A.W., Straub, A.C., Looft-Wilson, R., Johnstone, S.R., Araj, C.A., Best, A.K., Chekeni, F.B., Ravichandran, K.S., Penuela, S., et al. (2011). Pannexin1 regulates α1-adrenergic receptor-mediated vasoconstriction. Circ Res 109, 80–85.

Billaud, M., Sandilos, J.K., and Isakson, B.E. (2012). Pannexin 1 in the regulation of vascular tone. Trends Cardiovasc Med 22, 68–72.

Blum, A.E., Joseph, S.M., Przybylski, R.J., and Dubyak, G.R. (2008). Rho-family GTPases modulate Ca(2+) -dependent ATP release from astrocytes. Am J Physiol Cell Physiol 295, C231–241.

Boassa, D., Ambrosi, C., Qiu, F., Dahl, G., Gaietta, G., and Sosinsky, G. (2007). Pannexin1 channels contain a glycosylation site that targets the hexamer to the plasma membrane. J Biol Chem 282, 31733–31743.

Bruzzone, R., Hormuzdi, S.G., Barbe, M.T., Herb, A., and Monyer, H. (2003). Pannexins, a family of gap junction proteins expressed in brain. Proc Natl Acad Sci U S A 100, 13644–13649.

Burma, N.E., Bonin, R.P., Leduc-Pessah, H., Baimel, C., Cairncross, Z.F., Mousseau, M., Shankara, J.V., Stemkowski, P.L., Baimoukhametova, D., Bains, J.S., et al. (2017). Blocking microglial pannexin-1 channels alleviates morphine withdrawal in rodents. Nat Med 23, 355–360.

Chekeni, F.B., Elliott, M.R., Sandilos, J.K., Walk, S.F., Kinchen, J.M., Lazarowski, E.R., Armstrong, A.J., Penuela, S., Laird, D.W., Salvesen, G.S., et al. (2010). Pannexin 1 channels mediate ‘find-me’ signal release and membrane permeability during apoptosis. Nature 467, 863–867.

Chiu, Y.H., Jin, X., Medina, C.B., Leonhardt, S.A., Kiessling, V., Bennett, B.C., Shu, S., Tamm, L.K., Yeager, M., Ravichandran, K.S., et al. (2017). A quantized mechanism for activation of pannexin channels. Nat Commun 8, 14324.

Choi, W., Clemente, N., Sun, W., Du, J., and Lü, W. (2019). The structures and gating mechanism of human calcium homeostasis modulator 2. Nature 576, 163–167.

Dahl, G., and Keane, R.W. (2012). Pannexin: from discovery to bedside in 11±4 years? Brain Res 1487, 150–159.

Dahl, G., and Locovei, S. (2006). Pannexin: to gap or not to gap, is that a question? IUBMB Life 58, 409–419.

Deneka, D., Sawicka, M., Lam, A.K.M., Paulino, C., and Dutzler, R. (2018). Structure of a volume-regulated anion channel of the LRRC8 family. Nature. 558, 254–259

Deng, Z., He, Z., Maksaev, G., Bitter, R.M., Rau, M., Fitzpatrick, J.A.J., and Yuan, P. (2020). Cryo-EM structures of the ATP release channel pannexin 1. Nat Struct Mol Biol 27, 373–381.

Dolmatova, E., Spagnol, G., Boassa, D., Baum, J.R., Keith, K., Ambrosi, C., Kontaridis, M.I., Sorgen, P.L., Sosinsky, G.E., and Duffy, H.S. (2012). Cardiomyocyte ATP release through pannexin 1 aids in early fibroblast activation. Am J Physiol Heart Circ Physiol 303, H1208–1218.

Flores, J.A., Haddad, B.G., Dolan, K.A., Myers, J.B., Yoshioka, C.C., Copperman, J., Zuckerman, D.M., and Reichow, S.L. (2020). Connexin-46/50 in a dynamic lipid environment resolved by CryoEM at 1.9 A. Nat Commun 11, 4331.

Furlow, P.W., Zhang, S., Soong, T.D., Halberg, N., Goodarzi, H., Mangrum, C., Wu, Y.G., Elemento, O., and Tavazoie, S.F. (2015). Mechanosensitive pannexin-1 channels mediate microvascular metastatic cell survival. Nat Cell Biol 17, 943–952.

Gödecke, S., Roderigo, C., Rose, C.R., Rauch, B.H., Gödecke, A., and Schrader, J. (2012). Thrombin-induced ATP release from human umbilical vein endothelial cells. Am J Physiol Cell Physiol 302, C915–923.

Gulbransen, B.D., Bashashati, M., Hirota, S.A., Gui, X., Roberts, J.A., MacDonald, J.A., Muruve, D.A., McKay, D.M., Beck, P.L., Mawe, G.M., et al. (2012). Activation of neuronal P2X7 receptor-pannexin-1 mediates death of enteric neurons during colitis. Nat Med 18, 600–604.

Iglesias, R., Locovei, S., Roque, A., Alberto, A.P., Dahl, G., Spray, D.C., and Scemes, E. (2008). P2X7 receptor-Pannexin1 complex: pharmacology and signaling. Am J Physiol Cell Physiol 295, C752–760.

Jin, Q., Zhang, B., Zheng, X., Li, N., Xu, L., Xie, Y., Song, F., Bhat, E.A., Chen, Y., Gao, N., et al. (2020). Cryo-EM structures of human pannexin 1 channel. Cell Res 30, 449–451.

Kanneganti, T.D., Lamkanfi, M., Kim, Y.G., Chen, G., Park, J.H., Franchi, L., Vandenabeele, P., and Núñez, G. (2007). Pannexin-1-mediated recognition of bacterial molecules activates the cryopyrin inflammasome independent of Toll-like receptor signaling. Immunity 26, 433–443.

Karatas, H., Erdener, S.E., Gursoy-Ozdemir, Y., Lule, S., Eren-Koçak, E., Sen, Z.D., and Dalkara, T. (2013). Spreading depression triggers headache by activating neuronal Panx1 channels. Science 339, 1092–1095.

Kasuya, G., Nakane, T., Yokoyama, T., Jia, Y., Inoue, M., Watanabe, K., Nakamura, R., Nishizawa, T., Kusakizako, T., Tsutsumi, A., et al. (2018). Cryo-EM structures of the human volume-regulated anion channel LRRC8. Nat Struct Mol Biol 25, 797–804.

Kefauver, J.M., Saotome, K., Dubin, A.E., Pallesen, J., Cottrell, C.A., Cahalan, S.M., Qiu, Z., Hong, G., Crowley, C.S., Whitwam, T., et al. (2018). Structure of the human volume regulated anion channel. Elife 7.

Kern, D.M., Oh, S., Hite, R.K., and Brohawn, S.G. (2019). Cryo-EM structures of the DCPIB-inhibited volume-regulated anion channel LRRC8A in lipid nanodiscs. Elife 8.

Khan, A.K., Jagielnicki, M., McIntire, W.E., Purdy, M.D., Dharmarajan, V., Griffin, P.R., and Yeager, M. (2020). A Steric “Ball-and-Chain” Mechanism for pH-Mediated Regulation of Gap Junction Channels. Cell Rep 31, 107482.

Liao, M., Cao, E., Julius, D., and Cheng, Y. (2013). Structure of the TRPV1 ion channel determined by electron cryo-microscopy. Nature 504, 107–112.

Locovei, S., Bao, L., and Dahl, G. (2006a). Pannexin 1 in erythrocytes: function without a gap. Proc Natl Acad Sci U S A 103, 7655–7659.

Locovei, S., Wang, J., and Dahl, G. (2006b). Activation of pannexin 1 channels by ATP through P2Y receptors and by cytoplasmic calcium. FEBS Lett 580, 239–244.

Lu, P., Bai, X.C., Ma, D., Xie, T., Yan, C., Sun, L., Yang, G., Zhao, Y., Zhou, R., Scheres, S.H., et al. (2014). Three-dimensional structure of human γ-secretase. Nature 512, 166–170.

Ludtke, S.J., Baldwin, P.R., and Chiu, W. (1999). EMAN: semiautomated software for high-resolution single-particle reconstructions. J Struct Biol 128, 82–97.

Ma, W., Hui, H., Pelegrin, P., and Surprenant, A. (2009). Pharmacological characterization of pannexin-1 currents expressed in mammalian cells. J Pharmacol Exp Ther 328, 409–418.

Ma, Z., Siebert, A.P., Cheung, K.H., Lee, R.J., Johnson, B., Cohen, A.S., Vingtdeux, V., Marambaud, P., and Foskett, J.K. (2012). Calcium homeostasis modulator 1 (CALHM1) is the pore-forming subunit of an ion channel that mediates extracellular Ca2+ regulation of neuronal excitability. Proc Natl Acad Sci U S A 109, E1963–1971.

Maeda, S., Nakagawa, S., Suga, M., Yamashita, E., Oshima, A., Fujiyoshi, Y., and Tsukihara, T. (2009). Structure of the connexin 26 gap junction channel at 3.5 A resolution. Nature 458, 597–602.

Medina, C.B., Mehrotra, P., Arandjelovic, S., Perry, J.S.A., Guo, Y., Morioka, S., Barron, B., Walk, S.F., Ghesquiere, B., Krupnick, A.S., et al. (2020). Metabolites released from apoptotic cells act as tissue messengers. Nature 580, 130–135.

Mi, W., Li, Y., Yoon, S.H., Ernst, R.K., Walz, T., and Liao, M. (2017). Structural basis of MsbA-mediated lipopolysaccharide transport. Nature 549, 233–237.

Michalski, K., Syrjanen, J.L., Henze, E., Kumpf, J., Furukawa, H., and Kawate, T. (2020). The cryo-EM structure of a pannexin 1 reveals unique motifs for ion selection and inhibition. Elife 9.

Mou, L., Ke, M., Song, M., Shan, Y., Xiao, Q., Liu, Q., Li, J., Sun, K., Pu, L., Guo, L., et al.(2020). Structural basis for gating mechanism of Pannexin 1 channel. Cell Res 30, 452–454.

Myers, J.B., Haddad, B.G., O’Neill, S.E., Chorev, D.S., Yoshioka, C.C., Robinson, C.V., Zuckerman, D.M., and Reichow, S.L. (2018). Structure of native lens connexin 46/50 intercellular channels by cryo-EM. Nature 564, 372–377.

Oshima, A., Matsuzawa, T., Murata, K., Tani, K., and Fujiyoshi, Y. (2016a). Hexadecameric structure of an invertebrate gap junction channel. J Mol Biol 428, 1227–1236.

Oshima, A., Tani, K., and Fujiyoshi, Y. (2016b). Atomic structure of the innexin-6 gap junction channel determined by cryo-EM. Nat Commun 7, 13681.

Oshima, A., Tani, K., Hiroaki, Y., Fujiyoshi, Y., and Sosinsky, G.E. (2007). Three-dimensional structure of a human connexin26 gap junction channel reveals a plug in the vestibule. Proc Natl Acad Sci U S A 104, 10034–10039.

Panchin, Y., Kelmanson, I., Matz, M., Lukyanov, K., Usman, N., and Lukyanov, S. (2000). A ubiquitous family of putative gap junction molecules. Curr Biol 10, R473–474.

Pelegrin, P., and Surprenant, A. (2006). Pannexin-1 mediates large pore formation and interleukin-1beta release by the ATP-gated P2X7 receptor. EMBO J 25, 5071–5082.

Penuela, S., Bhalla, R., Gong, X.Q., Cowan, K.N., Celetti, S.J., Cowan, B.J., Bai, D., Shao, Q., and Laird, D.W. (2007). Pannexin 1 and pannexin 3 are glycoproteins that exhibit many distinct characteristics from the connexin family of gap junction proteins. J Cell Sci 120, 3772–3783.

Penuela, S., Gyenis, L., Ablack, A., Churko, J.M., Berger, A.C., Litchfield, D.W., Lewis, J.D., and Laird, D.W. (2012). Loss of pannexin 1 attenuates melanoma progression by reversion to a melanocytic phenotype. J Biol Chem 287, 29184–29193.

Pettersen, E.F., Goddard, T.D., Huang, C.C., Couch, G.S., Greenblatt, D.M., Meng, E.C., and Ferrin, T.E. (2004). UCSF Chimera--a visualization system for exploratory research and analysis. J Comput Chem 25, 1605–1612.

Poon, I.K., Chiu, Y.H., Armstrong, A.J., Kinchen, J.M., Juncadella, I.J., Bayliss, D.A., and Ravichandran, K.S. (2014). Unexpected link between an antibiotic, pannexin channels and apoptosis. Nature 507, 329–334.

Punjani, A., Rubinstein, J.L., Fleet, D.J., and Brubaker, M.A. (2017). cryoSPARC: algorithms for rapid unsupervised cryo-EM structure determination. Nat Methods 14, 290–296.

Qiu, F., Wang, J., Spray, D.C., Scemes, E., and Dahl, G. (2011). Two non-vesicular ATP release pathways in the mouse erythrocyte membrane. FEBS Lett 585, 3430–3435.

Qu, R., Dong, L., Zhang, J., Yu, X., Wang, L., and Zhu, S. (2020). Cryo-EM structure of human heptameric Pannexin 1 channel. Cell Res 30, 446–448.

Qu, Y., Misaghi, S., Newton, K., Gilmour, L.L., Louie, S., Cupp, J.E., Dubyak, G.R., Hackos, D., and Dixit, V.M. (2011). Pannexin-1 is required for ATP release during apoptosis but not for inflammasome activation. J Immunol 186, 6553–6561.

Ritchie, T.K., Grinkova, Y.V., Bayburt, T.H., Denisov, I.G., Zolnerciks, J.K., Atkins, W.M., and Sligar, S.G. (2009). Chapter 11 - Reconstitution of membrane proteins in phospholipid bilayer nanodiscs. Methods Enzymol 464, 211–231.

Ruan, Z., Orozco, I.J., Du, J., and Lu, W. (2020). Structures of human pannexin 1 reveal ion pathways and mechanism of gating. Nature 584, 646–651.

Sandilos, J.K., Chiu, Y.H., Chekeni, F.B., Armstrong, A.J., Walk, S.F., Ravichandran, K.S., and Bayliss, D.A. (2012). Pannexin 1, an ATP release channel, is activated by caspase cleavage of its pore-associated C-terminal autoinhibitory region. J Biol Chem 287, 11303–11311.

Santiago, M.F., Veliskova, J., Patel, N.K., Lutz, S.E., Caille, D., Charollais, A., Meda, P., and Scemes, E. (2011). Targeting pannexin1 improves seizure outcome. PLoS One 6, e25178.

Scemes, E., Spray, D.C., and Meda, P. (2009). Connexins, pannexins, innexins: novel roles of “hemi-channels”. Pflugers Arch 457, 1207–1226.

Scheres, S.H. (2012). RELION: implementation of a Bayesian approach to cryo-EM structure determination. J Struct Biol 180, 519–530.

Scheres, S.H. (2015). Semi-automated selection of cryo-EM particles in RELION-1.3. J Struct Biol 189, 114–122.

Seminario-Vidal, L., Kreda, S., Jones, L., O’Neal, W., Trejo, J., Boucher, R.C., and Lazarowski, E.R. (2009). Thrombin promotes release of ATP from lung epithelial cells through coordinated activation of rho- and Ca2+-dependent signaling pathways. J Biol Chem 284, 20638–20648.

Silverman, W.R., de Rivero Vaccari, J.P., Locovei, S., Qiu, F., Carlsson, S.K., Scemes, E., Keane, R.W., and Dahl, G. (2009). The pannexin 1 channel activates the inflammasome in neurons and astrocytes. J Biol Chem 284, 18143–18151.

Slotboom, D.J., Duurkens, R.H., Olieman, K., and Erkens, G.B. (2008). Static light scattering to characterize membrane proteins in detergent solution. Methods 46, 73–82.

Sosinsky, G.E., Boassa, D., Dermietzel, R., Duffy, H.S., Laird, D.W., MacVicar, B., Naus, C.C., Penuela, S., Scemes, E., Spray, D.C., et al. (2011). Pannexin channels are not gap junction hemichannels. Channels (Austin) 5, 193–197.

Strop, P., and Brunger, A.T. (2005). Refractive index-based determination of detergent concentration and its application to the study of membrane proteins. Protein Sci 14, 2207–2211.

Suadicani, S.O., Iglesias, R., Wang, J., Dahl, G., Spray, D.C., and Scemes, E. (2012). ATP signaling is deficient in cultured Pannexin1-null mouse astrocytes. Glia 60, 1106–1116.

Syrjanen, J.L., Michalski, K., Chou, T.H., Grant, T., Rao, S., Simorowski, N., Tucker, S.J., Grigorieff, N., and Furukawa, H. (2020a). Publisher Correction: Structure and assembly of calcium homeostasis modulator proteins. Nat Struct Mol Biol 27, 305.

Syrjanen, J.L., Michalski, K., Chou, T.H., Grant, T., Rao, S., Simorowski, N., Tucker, S.J., Grigorieff, N., and Furukawa, H. (2020b). Structure and assembly of calcium homeostasis modulator proteins. Nat Struct Mol Biol 27, 150–159.

Thompson, J.D., Higgins, D.G., and Gibson, T.J. (1994). CLUSTAL W: improving the sensitivity of progressive multiple sequence alignment through sequence weighting, position-specific gap penalties and weight matrix choice. Nucleic Acids Res 22, 4673–4680.

Thompson, R.J., Jackson, M.F., Olah, M.E., Rungta, R.L., Hines, D.J., Beazely, M.A., MacDonald, J.F., and MacVicar, B.A. (2008). Activation of pannexin-1 hemichannels augments aberrant bursting in the hippocampus. Science 322, 1555–1559.

Thompson, R.J., Zhou, N., and MacVicar, B.A. (2006). Ischemia opens neuronal gap junction hemichannels. Science 312, 924–927.

Tribet, C., Audebert, R., and Popot, J.L. (1996). Amphipols: polymers that keep membrane proteins soluble in aqueous solutions. Proc Natl Acad Sci U S A 93, 15047–15050.

Unger, V.M., Kumar, N.M., Gilula, N.B., and Yeager, M. (1999). Three-dimensional structure of a recombinant gap junction membrane channel. Science 283, 1176–1180.

Unwin, P.N., and Ennis, P.D. (1984). Two configurations of a channel-forming membrane protein. Nature 307, 609–613.

Wang, J., Ambrosi, C., Qiu, F., Jackson, D.G., Sosinsky, G., and Dahl, G. (2014). The membrane protein Pannexin1 forms two open-channel conformations depending on the mode of activation. Sci Signal 7, ra69.

Wang, J., Jackson, D.G., and Dahl, G. (2018). Cationic control of Panx1 channel function. Am J Physiol Cell Physiol. 315: C279–C289.

Yang, D., He, Y., Muñoz-Planillo, R., Liu, Q., and Núñez, G. (2015). Caspase-11 Requires the Pannexin-1 Channel and the Purinergic P2X7 Pore to Mediate Pyroptosis and Endotoxic Shock. Immunity 43, 923–932.

Zhang, K. (2016). Gctf: Real-time CTF determination and correction. J Struct Biol 193, 1–12.

Zheng, S.Q., Palovcak, E., Armache, J.P., Verba, K.A., Cheng, Y., and Agard, D.A. (2017). MotionCor2: anisotropic correction of beam-induced motion for improved cryo-electron microscopy. Nat Methods 14, 331–332.

